# Cysteine hyperoxidation rewires communication pathways in the nucleosome and destabilizes the dyad

**DOI:** 10.1101/2023.10.20.563272

**Authors:** Yasaman Karami, Emmanuelle Bignon

**Affiliations:** Université de Lorraine, CNRS, Inria, LORIA, F-54000 Nancy, France; Université de Lorraine and CNRS, UMR 7019 LPCT, F-54000 Nancy, France

## Abstract

Gene activity is tightly controlled by reversible chemical modifications called epigenetic marks, which are of various types and modulate gene accessibility without affecting the DNA sequence. Despite an increasing body of evidence demonstrating the role of oxidative-type modifications of histones in gene expression regulation, there remains a complete absence of structural data at the atomistic level to understand the molecular mechanisms behind their regulatory action. Owing to *μ*s time-scale MD simulations and protein communication networks analysis, we describe the impact of histone H3 hyperoxidation (i.e., S-sulfonylation) on the nucleosome dynamics. Our results reveal the atomic-scale details of the intrinsic structural networks within the canonical histone core and their perturbation by hyperoxidation of the histone H3 C110. We show that this modification involves local rearrangement of the communication networks and destabilizes the dyad, which could be important for nucleosome disassembly.

## INTRODUCTION

The regulatory effect of histone proteins post-translational modifications (PTM) onto DNA compaction and gene expression is a timely matter of research. If these modifications offer promising perspectives for the development of epigenetic therapies against a large panel of diseases, we are still far from understanding correctly their independent role on DNA compaction and their combinatorial effect. PTM consist in mostly reversible chemical modifications of amino acids that allow the regulation of proteins’ activity after their biosynthesis. There exists a broad spectrum of such modifications, and some of them (phosphorylation, glycosylation, methylation, acetylation) have been the privileged subject of investigations due to the robustness of their detection methods. At the molecular level, histone proteins (H3, H4, H2A, and H2B) assemble to form an octameric core around which *∼* 146 base pairs of DNA are wrapped, resulting in the first level of DNA compaction, the so-called the nucleosome core particle (NCP) – see Figure 1. Histone proteins share a common fold featuring three main *α* helices flanked by a C-term and/or N-term additional *α*helix (*α*C and/or *α*N). Nucleosome aggregation forms the chromatin, which undergoes dynamic structural exchanges between an open state (euchromatin) favoring DNA exposure for gene expression and a compacted state (heterochromatin) associated with gene silencing. Within this complex assembly, DNA compaction is regulated by a plethora of PTM, yet many aspects of the molecular mechanisms underlying this finely-tuned regulation remain to be unraveled.

**Figure 1.**
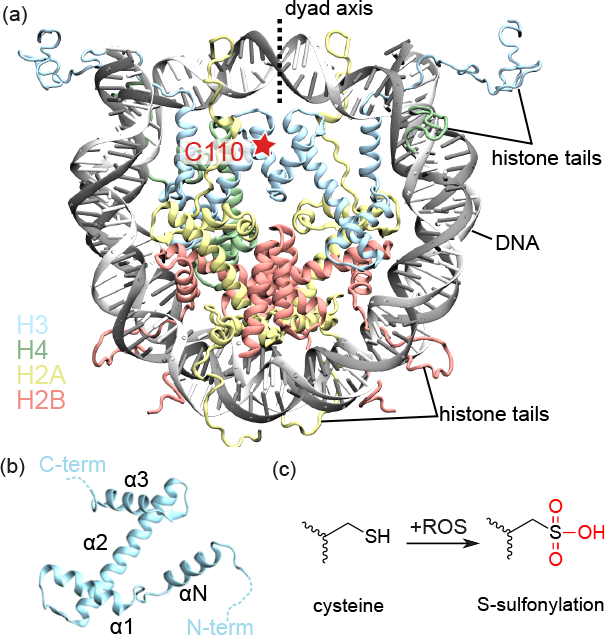
**(a)** Structure of a nucleosome modeled from the 1KX5 PDB (1) with symmetric histone H3 tails. Histone proteins are colored according to their type: H3 in blue, H4 in green, H2A in yellow, and H2B in orange. The position of the C110 residue on one histone H3 copy, that is targeted by oxidative PTM, is marked by a red star. **(b)** Canonical fold of the histone H3, featuring the conserved three main *α* helices and an additional *α*N helix in its N-terminal region. Other histone types can exhibit an additional *α*C as well, e.g., H2B. **(c)** Hyperoxidation of cysteines can induce its S-sulfonylation, which is an irreversible, deleterious oxidative PTM.

In this sense, structural biology and computational approaches are of major importance to get insights into the atomistic details underlying these phenomena. The first experimental structure of a nucleosome was published in 1997 by Luger et al.(2). Since then, technical improvements (e.g., the advent of cryoelectronic microscopy) led to the deposition of hundreds of NCP structures on the Protein Data Bank (PDB), gathering from mono- to 6-mer NCP assemblies(3) and large NCP-protein complexes. This accumulation of data also allowed to start unraveling the finely-tuned epigenetic regulation of chromatin dynamics by histone PTM and variants exchange, resorting to both experimental and theoretical approaches (4, 5). Yet, structures of NCP harboring PTM marks are rare, and a lot of aspects remain to be uncovered in order to get a better understanding of epigenetic regulation at the molecular level.

It is only recently that oxidative PTM (oxPTM) have gained interest from the biochemical community due to their importance in disease onset and for the development of redox-based therapies. Cysteines are preferential targets for oxidative modifications and act as redox switches in many proteins (6). They can react with nitric oxide (S-nitrosylation), glutathione (S-glutathionylation), hydrogen sulfide (S-persulfidation), and can undergo several other types of oxidation. Among the large spectrum of oxidatively-regulated proteins, nuclear proteins such as histones and proteins involved in DNA transcription, replication, and DNA damage repair have recently started to gather attention. Indeed, the activity of several partner proteins of the nucleosome can be regulated by oxPTM, such as histone de-acetylases (class 1 HDAC, sirtuin proteins…) (7, 8), transcription factors (STAT3, NF-*κ*B, AP-1…) (9, 10), or DNA repair enzymes (O6-alkylguanine-DNA-alkyltransferase, 8-oxoguanine glycosylase…) (11, 12), underlying the complexity and richness of oxidative stress and epigenetic processes crosstalk. The oxidation of histone proteins plays an important role in chromatin dynamics modulation, and oxidative stress-related alteration of gene expression is involved in aging and pathogenesis (e.g., cancer, neurodegenerative diseases, respiratory syndrome) (7). S-glutathionylation of the nucleosome has been shown to promote chromatin opening (13), while S-sulfonylation of histone H3 has been observed under oxidative stress (8). Yet, the importance of oxidative epiregulation of nucleosomal assemblies has been, until recently, widely neglected and their function and underlying molecular mechanisms remain to be uncovered.

Studies of histone core PTM location suggested that the ones located at the dyad (e.g., H4S47ph, H3K115ac, H3T118ph, H3K122ac) would favor the nucleosome disassembly, while those near the DNA entry/exit (e.g.,H3Y41ph, H3K56ac, H3S57ph) would promote DNA unwrapping without destabilizing the overall structure (14, 15, 16, 17, 18). Besides, computational investigations of the structural effect of H2A variants suggested allosteric effects and communication pathways involving the DNA double-helix (19). Importantly, the histone H3 cysteine (C110) is located near the dyad but is not in direct contact with the DNA helix, suggesting the its oxidative modification might impact the nucleosome dynamics in novel ways involving medium- to long-range effects that remain to be described.

Owing to molecular dynamics (MD) simulations and extensive structural post-analysis, we report here the first in-depth description of the nucleosome intrinsic protein structural networks, and its perturbation by histone H3 S-sulfonylation resulting from cysteine hyperoxidation. We show that this oxidative modification perturbs the DNA dynamics close in the dyad region. Such perturbation is induced by a rewiring of the local structural network, which results in the perturbation of the H3-H3 interface. We also scrutinize the DNA length and sequence effects onto the nucleosome protein structural network, by considering both *α*-satellite and 601 Widom DNA sequences. Our results provide unprecedented insights into the communication pathways within the histone core and their perturbation by an oxidative PTM, which sets the grounds for larger-scale mapping of molecular mechanisms underlying PTM regulation of the nucleosome dynamics at the atomic scale.

## MATERIALS AND METHODS

All simulations were performed using the NAMD3 software (20). System setup and analysis were performed using the AMBERTools20 suite of programs (21) and COMMA and Curves+ approaches (22, 23). The VMD 1.9.3 (24) and Pymol 2.5.5 (25) software were used for visualization and pictures rendering.

### System preparation

The *α*-satellite nucleosome starting structure for MD simulations was taken from Davey et al. crystal structure of the nucleosome featuring an *α*-satellite DNA sequence and histone tails (PDB ID 1KX5 (1)). Crystallographic waters and ions were removed. Force field parameters were taken from ff14SB (26), and bsc1 (27) and CUFIX (28) corrections were applied to improve DNA and the disordered tails description. The sulfonylated nucleosome starting system was created from the canonical nucleosome by mutating *in silico* the cysteine 110 of the first copy of histone H3. The modified residue name was set to OCS, consistently to what is found in the literature. Considering the very low pKa (*∼* 2) of sulfonated cysteines (29), the deprotonated form of OCS was used, with a total charge of -1. Parameters were generated for the sulfonylated cysteine using the antechamber protocol: i) the structure of the modified cysteine N- and C-term ends were capped by an acetyl (-OCH_3_) and a methylamino (-NHCH_3_) group, respectively; ii) this structure was optimized at the B3LYP/6-311+G** level and a frequency calculation was carried out to ensure that the energy reached a minimum; iii) Mertz-Kollman charges were computed at the HF/6-31+G* on the optimized structure; iv) the antechamber protocol was used to assign atom types and fit RESP charges; v) charges of the capping atoms were set to 0 and equally distributed on the other atoms to ensure a total charge of -1; vi) the AMBER library file was generated with the removal of capping atoms and the connectivity set onto the N and C atoms of the residue named OCS, using the tleap module of AMBER. The OCS parameters and MD input files are available in SI. Each system (control and sulfonylated) was soaked into a TIP3P truncated octahedral water box applying a 20Å buffer between the nucleosome and the edges of the box. A 0.150 M salt concentration was ensured by randomly adding 378 Na^+^ and 233 Cl^*−*^ ions. The final systems gathered a total of *∼*400,000 atoms including *∼*123,000 water molecules.

### Molecular dynamics simulations

Each system was first subjected to a 30,000-step energy minimization using the conjugate gradient algorithm. Then, four subsequent equilibration runs of 10 ns were carried out at 300K with decreasing constraints on the biomolecule’s atoms. The time step was then increased from 2 fs to 4 fs by using the Hydrogen Mass Repartitioning algorithm (30) in addition to the SHAKE and RATTLE ones (31), and a 2 *μ*s production run was performed in the NPT ensemble. The temperature and the pressure were kept constant using the Langevin thermostat with a 1 ps^*−*1^ collision frequency and a Langevin piston barostat with a damping time scale of 50 fs and an oscillation period of 100 fs. Electrostatics were treated using the Particle Mesh Ewald approach (32) with a 9 Å cutoff.

For each system, six replicates were carried out with random initial velocities in order to ensure the statistical significance of the results, resulting in a total of 24*μ*s of simulation time. Velocities, box information and coordinates were recorded every 25,000 steps (i.e., every 0.5 ns and 1 ns for the equilibration and production runs, respectively).

MD ensembles of the 601 Widom nucleosome were taken from a previous study from us and collaborators (33), which consisted in three replicates amounting for a total of *∼* 15 *μ*s performed in the same conditions as the present study.

### Structural analysis

#### Protein network analysis

For each system, the network of pathways and communication blocks were identified using COMMA2 (22). In COMMA2, communication pathways are chains of residues that are not adjacent along the sequence, are linked by non-covalent interactions (hydrogen bonds or hydrophobic contacts) and communicate efficiently. Communication efficiency or propensity is expressed as (34):

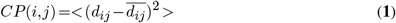

where *d*_*ij*_ is the distance between the C*α* atoms of residues *i* and *j* and *d*_*ij*_ is the mean value computed over the entire set of conformations. These pathways form the protein communication network (*PCN*), in which nodes correspond to the residues of the protein and edges connect residues adjacent in a pathway. COMMA2 extracts connected components from the graph by using depth-first search algorithm to identify the protein dynamical units. These units are referred to as “communication blocks” (see (34) for detailed descriptions).

Hydrogen bonds networks were calculated using the the HBPLUS algorithm (35). This algorithm detects hydrogen bonds between the donor (D) and acceptor (A) atoms using the following geometric criteria: *(i)* maximum distances of 3.9 *Å* for D-A and 2.5 *Å* for H-A, *(ii)* minimum value of 90% for D-H-A, H-AAA and D-A-AA angles, where AA is the acceptor antecedent. For every pair of residues, we assigned an interaction strengths as the percentage of conformations in which a hydrogen bond is formed between any atoms of same pair of residues. We then merged the results from all replicates of each system. We reported hydrogen bonds that are present for more than 40% of the simulation time (with strength values greater than or equal to 0.4). Only the proteins were taken into account for the COMMA2 analysis).

#### Per residue flexibility contribution

Per residue contribution to the overall flexibility of the system was calculated for DNA and histones, using a PCA-based machine learning script that was successfully used on similar DNA-protein and nucleosomal systems in previous studies (36, 37, 38). In this script, the internal coordinates of the residues are extracted from the MD ensembles, and a covariance matrix is generated, which eigenmodes and eigenvectors respectively represent the system’s modes of motion and their amplitude. The main fluctuations of the system are encrypted in the highest amplitude eigenmodes, and per residue normalized contribution to these modes of motion are calculated that translates their contribution to the overall flexibility of the system.

This analysis has been conducted onto the control and S-sulfonylated systems, taking into account the six MD simulations replicates performed for each system. Calculations for the DNA helix and the protein were performed separately, and the histone tails were not included to prevent statistical noise.

#### Other structural descriptors

The cpptraj AMBER module was used to calculate all distances and RMSD values, and to perform clustering analysis. For the monitoring of hydrogen bonds/salt bridges distances involving lysines, arginines, aspartates or glutamates side chains, the NZ, CZ (or NH2/NH1 if relevant), CG, and CD atoms were respectively taken into account for the distance calculation. This allowed to avoid monitoring any jumps that could arise from the rotation of the terminal charged moiety. The Curves+ software (23) was used to monitor the DNA structural parameters around the dyad section (between superhelical locations SHL-1 and SHL+1). The structure of the 21-bp section around SHL0 was extracted from the MD ensemble with 1 frame per ns. These structures were submitted to Curves+ analysis and the corresponding statistical values were computed from the results.

## RESULTS

For sake of clarity, the two sides of the nucleosome are referred to as face A and face B, and the amino acids of the second copy of each histone are named with an extra apostrophe (e.g., R129’).

### Communication blocks and pathways in the native nucleosome

#### Communication blocks

The protein structural network analysis of the *α*-satellite nucleosome reveals fifteen communication blocks within the histone core, i.e. fifteen independent groups of residues that mediate short- and long-range communication pathways within the overall structure - see Figure 2-a. The arrangement of these blocks on the histone core is mostly symmetric, with seven blocks for each face (A and B) of the nucleosome.

**Figure 2.**
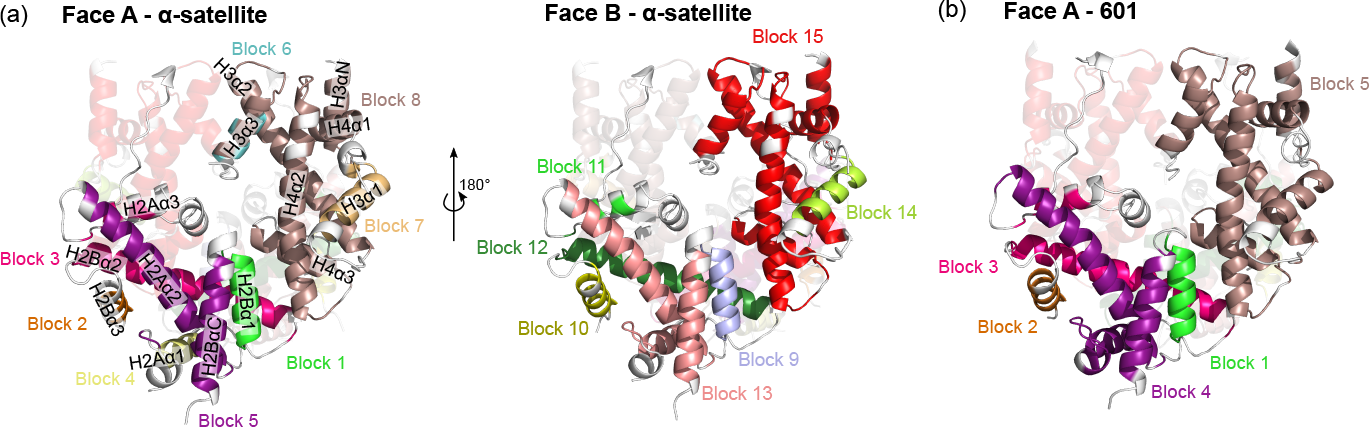
**(a)** Communication blocks within the canonical nucleosome featuring the *α*-satellite DNA sequence. Due to the intrinsic symmetry of the histone core, the blocks on face A (left) and B (right) of the nucleosome are mostly equivalent. Histone helices are labeled on Face A. **(b)** Communication blocks with the 601 Widom DNA sequence. Only face A is shown, see Figure S1 for face B. Histone tails are not displayed.

These communication blocks are distributed along the H2A-H2B and H3-H4 dimers but do not spread to wider architectures. Interestingly, the H2A-H2B dimers are divided in five blocks whereas the H3-H4 dimers exhibit a more compact structure with two main blocks. This could be linked to the fact that the nucleosome is built by the docking of two separate H2A-H2B dimers onto a (H3-H4)_2_ tetramer (39).

In each H2A-H2B dimer, the two main blocks involve residues of both H2A and H2B which might be important for the dimer formation. On the first face of the nucleosome (face A), the biggest block (block 5) involves H2B *α*C helix and H2A *α*2 helix. The second biggest (block 3) gathers H2B *α*2 helix and H2A *α*3 helix. The three other blocks are located on H2B *α*1 (block 2) and *α*3 (block 1), and H2A *α*3 (block 4). Noteworthy, a slight asymmetry is observed on the other face of the nucleosome (face B), in which a larger block (block 13) spreads onto H2B *α*C and H2A *α*2/*α*3, with the block gathering H2B *α*2 helix and H2A *α*3 helix (block 3 on face A) split in two (blocks 11 and 12).

The H3-H4 dimers exhibit more cohesive blocks. On both faces of the nucleosome, a large block involves the three H4 main helices and H3 *α*N/*α*2/*α*3 (Block 8/15 on face A/B, respectively), and H3 *α*1 forms an isolated block (block 7/14). On face A a small block is found in H3 *α*3 helix, intertwined with the major one, that involves only four amino acids (I111’, L113’, R115’, and I117’). A detailed list of the number of residues in each block is given in Table S1.

Taking advantage of 3×5 *μ*s MD ensembles of the 601 Widom sequence nucleosome from a previous work (33), we performed the same analysis to get insights into DNA sequence effects onto the histone core communication blocks and pathways. The 601 structure exhibits less (9 instead of 15) but larger communication blocks - see Figure 2-b. This suggests existence of longer range communication pathways in the 601 Widom system compared to the *α*-satellite one. H2A *α*3 here forms a large block with H2B *α*C and H2A *α*2 (block 4, instead of two separated blocks with the *α*-satellite sequence), and the H3-H4 is involved in a compact isolated block (block 5). The same trend is observed on face B of the 601 nucleosome, yet it shows even larger communicating regions by the fusion of blocks 1 and 3 - see Figure S1. This tighter communication network could come from the stronger packing of the DNA around the histones, that could sterically reinforce the intrinsic interactions within the histone core by limiting its dynamical behavior.

#### Communication pathways

In order to have a better understanding of the wiring of the communication blocks, communication pathways within the *α*-satellite nucleosome structure were scrutinized. The analysis of the communication pathways shed light on a large communication network in which communication hubs could be pinpointed, i.e. the amino acids that might be crucial for the system’s structural stability and allosteric regulation - see Figure 3-a. Very interestingly, the major hubs in the nucleosome structure are found on histone H3 *α*2 helix - the full list of number of paths per residue is given in SI. In both histone H3 copies, the M90-T107 section exhibits the larger hubs in the entire structure. As this helix is involved in the large H3-H4 communication block, it acts as a pivot for long range interactions within this dimer. At the one end, it mediates many pathways in the direction of H3 *α*1 and H4 *α*1 helix (itself highly connected to H3 *α*N and H4 *α*2) - see Figure 3-b. Of note, the C110 is located next to T107, that appears to be a dense hub, suggesting that the modification of this cysteine might have a crucial impact on the communication pathways that depend on the H3 *α*2 helix. Indeed, it has been suggested that residues adjacent to communication hubs can as well impact the structure and function of proteins (40). At the other end of H3 *α*2 helix, hubs are involved in few connections with H4 *α*1 again, and H3 *α*3 - see Figure 3-c. As it is involved in several communication pathways, H4 *α*1 also harbors quite populated hubs, from K31 to V43.

**Figure 3.**
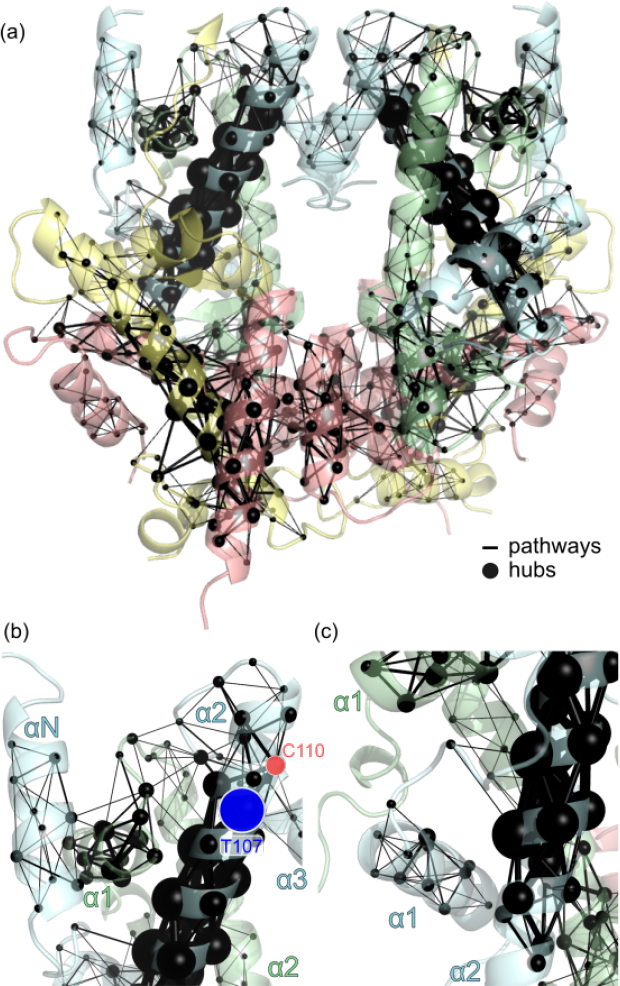
**(a)** Mapping of communication pathways (black lines) and hubs (black spheres) in the *α*-satellite nucleosome. The larger the line/sphere, the more populated the pathway/hub. **(b)** Pathways around the top of the H3 *α*2 helix, connecting H3 *α*3 and H4 *α*1. C110 and T107 are displayed as red and blue circles, respectively. **(c)** Pathways at the bottom of the H3 *α*2 helix, showing few connections with H3 *α*3

As already seen in the description of the communication blocks, pathways in the H2A-H2B dimers are much more fragmented than in H3-H4 - see Figure S2. Hubs can be pinpointed in H2A *α*2 (residues L51 to L63) and to a lesser extent in H2B *α*C (residues A107 to Y118), mostly responsible for communication blocks 5 and 13 identified previously. Less populated hubs are also observed in H2B *α*2, mostly from A55 to V66. Overall, H2A-H2B hubs appear much less dense than hubs in the H3-H4 dimers, which also explains why communication blocks in H2A-H2B are more fragmented.

Pathways analysis in the 601 nucleosome resulted in the same trends as for the *α*-satellite system. The only difference one can observe is a higher density of communication pathways between the histone helices, which translates into the more compact communication blocks as observed above - see Figure S1. However, the density of the communication hubs remains the same, with the H3 *α*2 helix exhibiting the highest contribution to the overall pathways.

### S-sulfonylation reshapes the H3-H3 contacts by rewiring the local interaction network

#### Local perturbation of the H3-H3 contacts

In the nucleosome structure, the bundle formed by H3 and H3’ *α*2 and *α*3 helices is of crucial importance for the (H3-H4)_2_ tetramer assembly. H3C110 hyperoxidation into its sulfonylated counterpart (OCS) induces a rewiring of the local interaction network at the H3-H3 interface, perturbing the stable canonical architecture.

The most pronounced structural change is induced by the rotation of H3R129’ side chain to form a hydrogen bond with OCS, which provokes the displacement of H3 and H3’ *α*2 helices ends - see Figure 4. The representative structures and corresponding distance values are showed only for the first MD replicate, as this phenomena is observed in all the MD replicates - see Figure S3. At the beginning of the simulation, the structure of the nucleosome bearing OCS is very close to the unmodified one (in purple and transparent blue, respectively in Figure 4-a). However, a conformational change is observed after *∼*100 ns of simulation, with the rotation of R129’ inwards to interact with OCS, that pushes H3 and H3’ *α*2 helices away from each other for the rest of the simulation - see Figure 4-b. Interestingly, the *α*3 helix position is very weakly perturbed by these changes, as suggested by a very modest drift of the CA backbone atom position of R129’ (1.7 *±* 0.4 Å for MD1, 1.8 *±* 0.6 Å over all MD replicates) compared to its CZ side chain atom (5.6 *±* 1.0 Å for MD1, 5.0 *±* 1.5 Å over all MD replicates), as showed in Figure 4-c. On the contrary, a pronounced drift of OCS CA atom with respect to its initial position is observed (3.2 *±* 0.7 Å for MD1, 2.7 *±* 1.1 Å over all MD replicates), *± ∼* which illustrates the displacement of H3 *α*2 helix end. The evolution of the distances between the center of mass of the H3 and H3’ *α*2 helices ends (taken as residues 105-115 and 105’-115’) also shows that these helices are moving away from each others, with an overall increase of 3.5 Å and an average distance of 16.1 0.8 Å compared to a value of 12.5 *±* 0.2 Å in the control simulations - Figure 4-d. Interestingly, while H3R129’ is initially close enough to H3E105 to form a salt bridge, this interaction does not hold upon the presence of OCS, with which R129’ interacts preferentially as illustrated by the increase of the distance between R129’ CZ and E105 CD atoms (from *∼* 7 Å to *∼* 13 Å, average of 12.4 *±* 1.4 *∼ ∼ ±* Å) and the decrease of the distance between R129’ NH2 and OCS SG atoms (from 13 Å to 4 Å, average of 4.6 2.0 Å).

**Figure 4.**
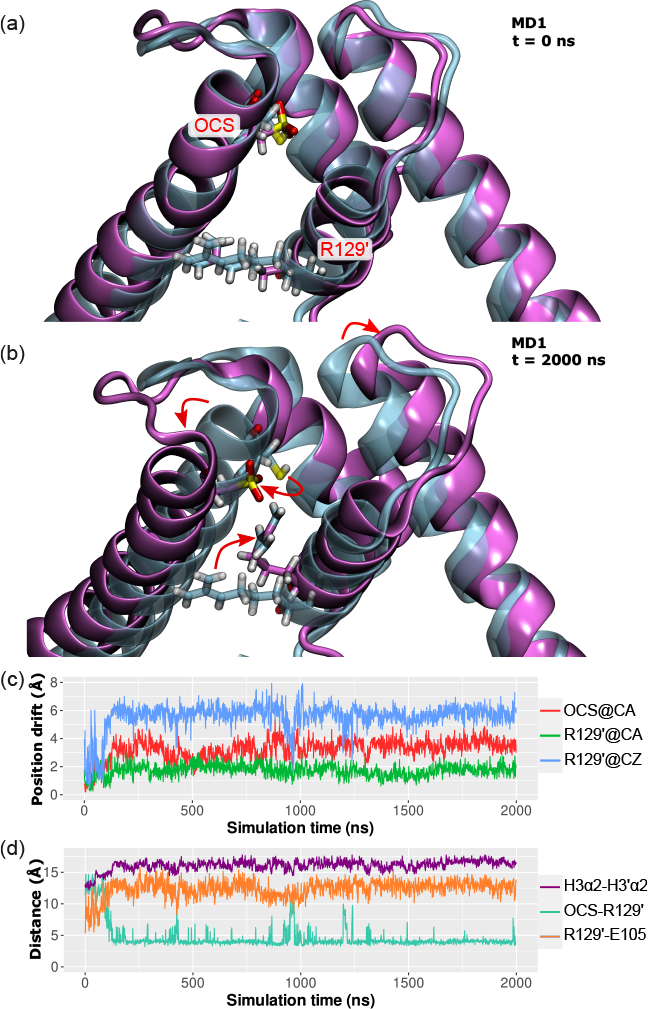
Superposition of the S-sulfonylated structure (purple) at **(a)** t = 0 ns and **(b)** at t = 2000 ns of the MD1 replicate onto the unmodified crystal structure (transparent cyan). Both copies of histone H3 are shown. The OCS sulfonylated residue and R129’ on the facing copy are displayed in licorice to illustrate their interaction upon sulfonylation. **(c)** Positional drift (in Å) of OCS CA atom (red) and R129’ CA (green) and CZ (blue) atoms along the MD1 replicate. It is calculated along the simulation as the distance of each atom to its initial position in the crystal structure. **(d)** Evolution of the distances between the ends of the histones H3 *α*2 helices (taken as the residues 105 to 115, in purple), between OCS SG atom and R129’ NH2 atom (cyan), and between R129’ CZ atom and H3E105 CD atom (orange).

The displacement of H3 and H3’ helices also involves some changes in the canonical interaction patterns of the four-helix bundle. One very stable symmetrical hydrogen bond in the non-modified nucleosome simulations is found between H113 and D123 on the facing H3 copy, with an average distance between H113 HE2 and D123 CG atoms of 3.3 *±* 0.3 Å for both nucleosome sides - see Figures S4 and S5. In the simulations with OCS these interactions are destabilized, especially for the one with H113 on the same H3 copy as OCS (i.e., H113-D123’). Average distances are in this case increased to 10.7 *±* 3.2 Å (H113-D123’) and 4.3 *±* 1.2 Å (H113’-D123). The presence of OCS might induce steric and electrostatic hindrance that prevents the formation of the canonical hydrogen bond network. The mapping of the hydrogen bond networks perturbation confirms the aforementioned loss and gain of interactions, and also revealing longer-range impact on the interaction between histones H3 C-termini and the H2A-H2B dimers - see Figure S6.

#### Rewiring of the communication blocks and pathways

The perturbation of the local interactions network near OCS results in a re-shaping of the overall communication blocks - see Figure 5-a. Noteworthy, the H2A-H2B dimers exhibit bigger blocks than in the unmodified system. On face A, three blocks can be distinguished that independently gather H2A *α*3 and H2B *α*1-2 (Block 1), H2B *α*3 (Block 2), and H2A *α*1-2 and H2B *α*N (Block 3). On face B, only two blocks are found, with the H2B *α*3 still isolated in Block 5 and the rest of the H2A- H2B *α* helices all gathered in Block 6. Besides, H3-H4 dimers are deferentially impacted, rather it is on OCS side or not. On face A, a single block is observed (Block 4), whereas on face B H3 *α*3 does not communicate with the other helices and H3 *α*1 is isolated in Block 8. The other helices are gathered in the large Block 9. Interestingly enough, the OCS residue is located in Block 9, while the H3 *α*3 helix that faces it is disconnected from the rest of the dimer, with only a small communication block (Block 7) of four amino acids (Q125, A127, R129, R131).

**Figure 5.**
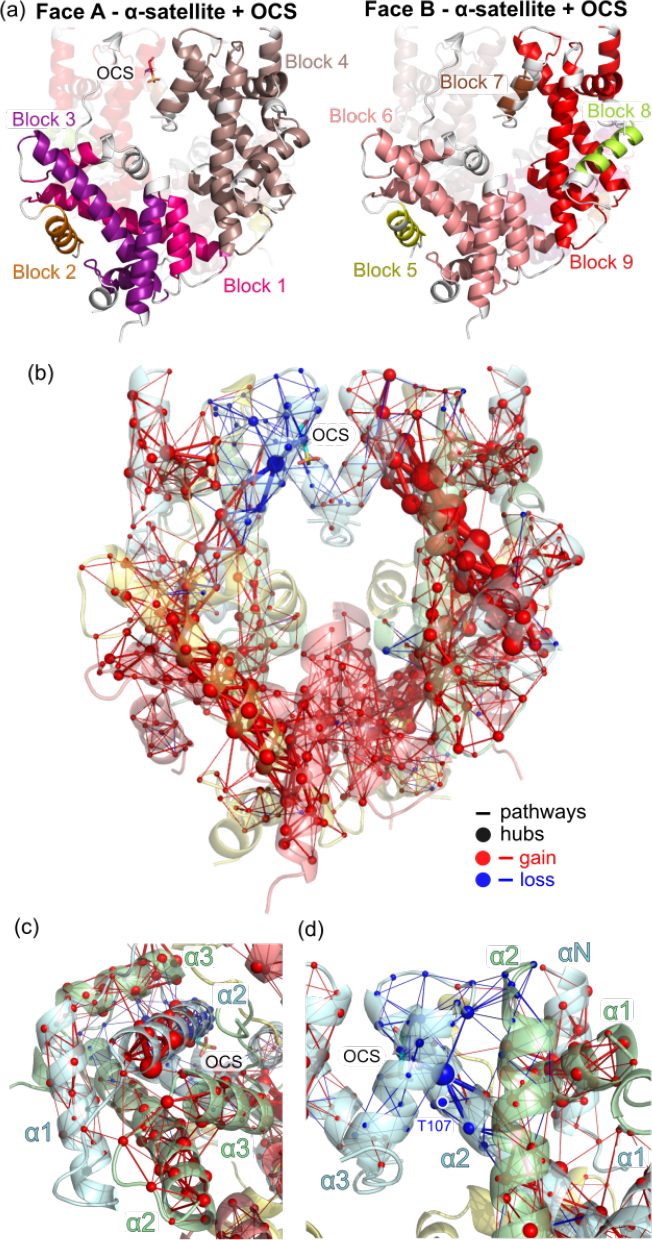
**(a)** Communications blocks in the *α*-satellite nucleosome with cysteine S-sulfonylation (OCS), on face A and face B. **(b)** Perturbation of communication pathways upon S-sulfonylation, viewed from face B. Red paths/hubs are gains with the modification, blue are losses. **(c)** Bottom view of H3’ *α*2 helix (in communication Block 4), through which multiple pathways connect other H3’ and H4’ helices. **(d)** Weakened pathways around the S-sulfonylated cysteine. T107 is highlighted by a white-blue circle.

A more in-depth look at the communication pathways allows to gain insights into the molecular mechanisms behind the blocks reorganization. The overall map of the pathways reveals an increase of the hubs importance in H3 and even H2A *α*2 helices - see Figure 5-b. The same H2A *α*2 hubs are found compared to the canonical nucleosome, but they funnel an increased number of inter-helices pathways within H2A-H2B, explaining the presence of larger communication blocks in these dimers. Of note, both H2A-H2B dimers show enhanced communication pathways, yet this trend is even more pronounced for face B than face A, which also rationalize the asymmetry of the blocks - see Figure S2.

The major hubs in H3 *α*2, already observed in the unmodified nucleosome, are also found to be more populated upon S-sulfonylation. At the bottom of this helix, the communication network between H3 *α*1, and H4 *α*1-3 shows much more connections - see Figure 5-c. At its top, an asymmetry between H3 (bearing OCS) and H3’ is observed, as one could expect. On H3’, communication pathways are found similar to the canonical nucleosome, yet the hubs importance appear to be slightly increased. On H3, where OCS perturbs the interaction network as above-described, there is a drastic loss of connection between H3 *α*2 and *α*3 helices, which decreases the importance of the neighbouring hubs such as T107 - 5-d. Weak pathways are found in H3 *α*3 but they remain isolated, with no connection to the other helices. The hydrogen bond network analysis shows a reorganization of the interactions around the OCS residue, and a loss of contacts between the H3/H3’ C-termini and H2A and H4 - see Figure S6.

### S-sulfonylation destabilizes DNA near the dyad

#### Increased DNA flexibility at SHL0/SHL-1

Besides the extensive description of histone core communication networks and their rewiring by S-sulfonylation, the impact of this modification on the DNA helix structure and dynamics was also assessed. The structural descriptor that was the most impacted by histone H3 S-sulfonylation is the flexibility of the nucleic acids in the dyad region. The per residue contribution to the overall DNA flexibility was computed for the S-sulfonylated system and the unmodified one. The mapping of the deviation of these contributions upon S-sulfonylation reveals an increase of flexibility not only at the DNA entry/exit sites but also, more interestingly, near the dyad (SHL0) - see Figure 6-a. In the nucleosome structure, the dyad region is the most stable part of nucleosomal DNA, and DNA-protein interactions near this axis have a crucial role for nucleosome stability and the regulation of its assembly/disassembly (14, 17, 18). The hyperoxidation of histone H3 C110 results in an asymmetric destabilization of the dyad region, with a more pronounced effect on the SHL0/SHL-1 region opposite to the PTM site (i.e., not directly above the PTM). The DNA entry/exit are also destabilized by the presence of OCS, with a much more pronounced effect on the side of the nucleosome opposite to OCS - Face B, see 6-b. However, this only concerns less than 10 base-pairs (bp) sections on DNA termini. Noteworthy, weak deviations are also observed on longer-range distances: a very localized stiffening of nucleotides (dA174 and dA175 at SHL-4.5, dC237 and dT238 at SHL1.5), and a slight destabilization of strand1 at SHL4/4.5.

**Figure 6.**
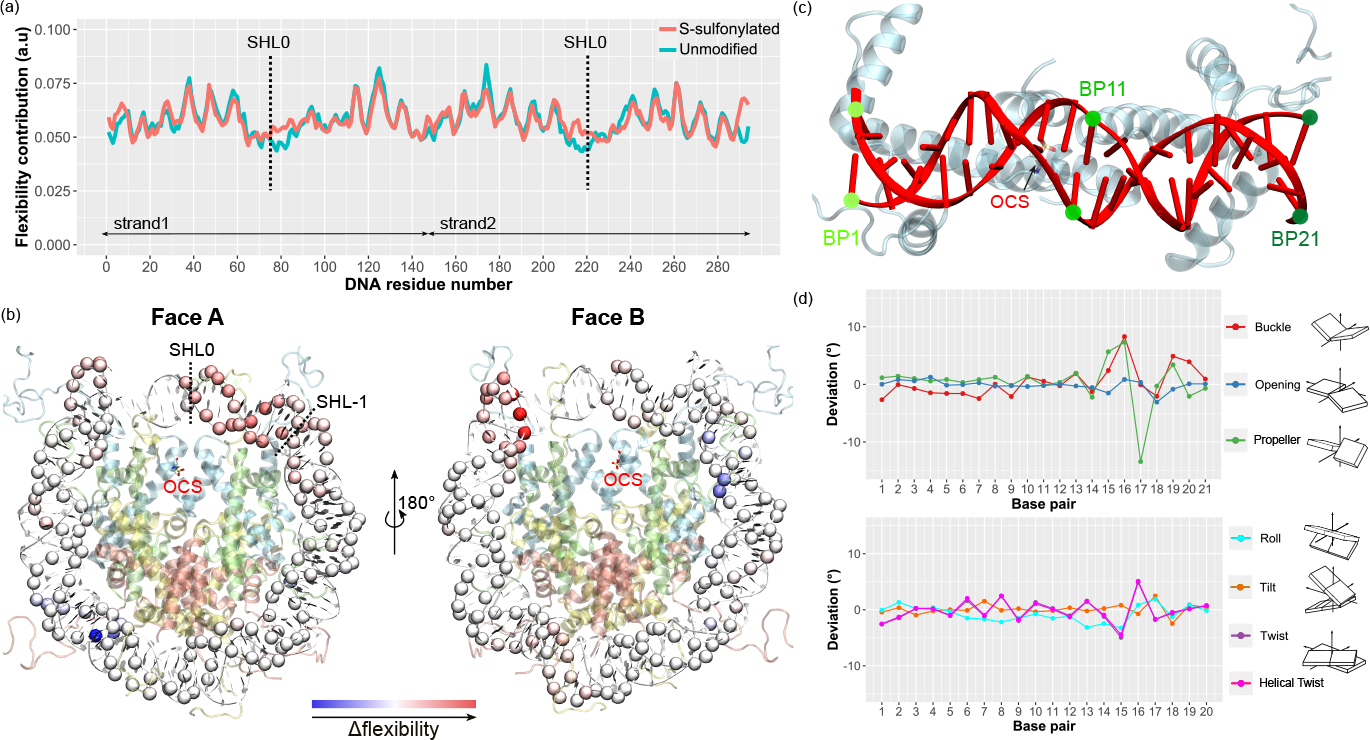
**(a)** Per residue flexibility contribution of the two DNA strands. SHL0 is indicated as it is, besides the DNA entry/exit, the region the most perturbed by OCS. **(b)** Projection of the perturbation of the per residue flexibility in DNA upon S-sulfonylation, with respect to the unmodified system. Red: higher flexibility, blue: lower flexibility, white: no perturbation. The S-sulfonylated cysteine is displayed in licorice and superhelical locations SHL0 and SHL-1 are indicated on Face A to highlight the enhanced flexibility in this region. **(c)** Top view of the 21-bp DNA section taken for the structural parameters calculation (i.e., between SHL+1 and SHL-1). Base pairs 1, 11, and 21 are labeled in green. Histones H3 and OCS are displayed in transparent. The remaining of the system is not shown. **(d)** Deviation of the intra- (top) and inter- (bottom) base pair angular parameters of the 21-bp DNA at the dyad in the S-sulfonylated nucleosome with respect to the unmodified system. Schematic representations of the parameters are provided, with each nucleobase displayed as a square. Inter-base pair parameters are calculated by couples of adjacent base pairs (e.g., bp1-bp2, bp2-bp3).

In order to probe the molecular mechanisms driving the OCS-induced DNA destabilization near the dyad axis, an in-depth structural analysis of the 21-bp section around the dyad axis was performed, and DNA-protein interactions were monitored.

#### DNA structural parameters at the dyad

Structural descriptors of the 21-bp DNA section around the dyad were computed on the MD ensembles with and without S-sulfonylation - see Figure 6-c and Figures S7 and S8. The OCS-induced deviation of intra-base pair, inter-base pair, and base pair axis parameters did not show any drastic deformation of the DNA double helix, but interestingly the largest perturbations are localized in the second half of the 21-bp section, i.e., in the SHL0/SHL-1 region that was found to experience an enhanced flexibility. For instance, the buckle and propeller angles deviations is more pronounced within base pairs 15- 19, corresponding to residues 78-82 and 213-217 on strands 1 and 2, respectively - see Figure 6-d. Likewise, the inter-base pair twist angles exhibit peaks in this region. Overall, S-sulfonylation seems to only weakly perturb the DNA structure at the dyad, yet the largest deviations of the structural parameters with respect to the unmodified system are found in the SH0/SHL-1 region for which the flexibility was increased with the presence of OCS.

#### Interactions between DNA and the histone core lateral surface

Canonical interactions between DNA and the histone residues at the lateral surface of the octamer were listed from analysis of the *α*-satellite nucleosome experimental structure and from the literature, and monitored in the simulations of the unmodified and S-sulfonylated *α*-satellite nucleosomes - see Table S2 and Figures S9 and S10. While most of the hydrogen bonds and salt bridges were unchanged by the hyperoxidation of histone H3, some of them located near the dyad exhibited a larger deviations in the S-sulfonylated system. It is the case for interactions involving H3K115 and H3T118, showing a broader distribution with OCS than in the unmodified system. On histone H3/H3’, K115 NZ atom distance to dA220 phosphate is of 7.0 *±* 2.0 Å with OCS and 6.1 *±* 1.1 Å without, and K115’ NZ atom distance to dA73 phosphate is of 7.2 *±* 1.8 Å with OCS and 6.1 *±* 1.2 Å without. Interestingly, the deviation of the distance between T118’ HG1 and dG218 phosphate is larger with OCS (3.2 *±* 0.9 Å with OCS vs 2.9 *±* 0.2 Å without), which could be a result of the H3 helices displacement aforementioned, participating to the increase of DNA flexibility in this region. Interestingly, the hydrogen bond network mapping showed an enhanced interaction between H3T118’ and H4R45’, which might weaken their contact with other partners including DNA - see Figure S6. Of note, the flexibility analysis of the protein dimers did not show any drastic deviation upon S-sulfonylation in the structured parts of the histone core, yet the H3 *α*3 helix facing the OCS site exhibits a slight increase with is not observed in its H3’ non-modified counterpart - see Figure S11.

## DISCUSSION

Owing to extensive all-atom molecular dynamics simulations and in-depth structural analysis, we described the communication networks within the histone core of the canonical nucleosome with both *α*-satellite and 601 Widom sequence. We also investigated the perturbation upon hyperoxidation of histone H3C110, showing that the S-sulfonylation PTM of this residue results in the destabilization of the DNA at the dyad and a rewiring of the interactions network within the histone core.

The detailed map of the communication pathways within the canonical nucleosome offers a novel way to apprehend how histone proteins synergetically interact with one another. Our results show that communication blocks are localized in separate H2A-H2B or H3-H4 units, i.e., no inter-dimer communication pathways are observed - see Figures 2 and S1. The H2A-H2B dimers is more fragmented, which underlines a less compact interaction network than in their H3-H4 counterparts. Despite their more intense intrinsic communication networks H3-H4 dimers do not share any communication pathway, which could explain why H3-H4 dimers are found to be very stable while the (H3-H4)_2_ tetramer remains less stable than H2A-H2B (41). Interestingly, while only very weak differences are observed with the 601 Widom sequence compared to the *α*-satellite nucleosome, larger communication blocks are found within the dimers, suggesting a more compact architecture which might result from the tighter packing of DNA (146-bp vs 147-bp in the *α*-satellite sequence) - see Figure 2-a and S1.

The in-depth analysis of communication pathways allowed to pinpoint key-residues acting as communication hubs within the nucleosome structure, for both 601 and *α*- satellite sequences. The most connected hubs are found in histone H3 *α*2 helix (M90 to T107), which funnel extensive communication pathways within the H3-H4 dimer - see Figure 3. None of them undergo PTM, except T107 that can be phosphorylated (42). Besides, as C110 is directly connected to T107, suggesting that its modification might have important consequences on the H3-H4 communication network as detailed below for S-sulfonylation. Likewise, only few residues in H4 *α*1 and H2A *α*2 largest hubs are known to be PTM sites (H4K31 and H2AT59). As was already observed for point mutations (40), PTM sites are important to regulate the system’s structure, but they are not necessarily large communication hubs and instead are most frequently located in their vicinity. Interestingly enough, few of these hubs are also mutational hotspots in several types of cancer (H3E97, H3E105, H2AE56) (43).

Noteworthy, the communication pathways are exclusively found in the structured histone core, not on the tails. As PTM on the histone tails are mostly involved in partner protein-mediated regulation of the nucleosome architecture or direct modulation of DNA-tails interactions, PTM in the histone core might have a different role with a direct influence onto DNA-histone interactions and nucleosome dynamics (44). It is important to underline that the COMMA2 analysis of the communication networks does not include the DNA helix, yet communication pathways might also transit through it via histone-DNA interactions as previously suggested by Bowerman and Wereszczynski (19). These results nevertheless provide an in-depth description of the histone units cooperativity, and the next step will be to improve COMMA2 towards the inclusion of DNA in the network analysis.

Histone H3 are the only units bearing cysteine residues, at position 110 in H3.2 and at positions 96 and 110 in variants H3.1 and H3.3. C110 is located in the four-helix bundle at the H3-H3 interface and is known to undergo diverse types of oxidative PTM (7). Our simulations brought insights into the structural impact of S-sulfonylation, a PTM resulting from cysteine hyperoxidation, which revealed that this modification impacts not only the histone core intrinsic communication pathways, but also destabilizes the DNA structure near the dyad axis. Generally, PTM located on the lateral surface of the histone core and in contact with the dyad DNA (H4S47ph, H3K115ac, H3T118ph, H3K122ac) are supposed to promote the nucleosome disassembly by directly impairing histone-DNA interactions (16, 18). In the case of H3C110 S-sulfonylation, the structural effect is not direct, because C110 side chain is located *∼*15 Å away from any nucleic acid. Instead, this modification induces a reorganization of the interaction network, stabilizing the four-helix bundle at the H3-H3 interface, resulting in the displacement of the H3 *α*2 helices - see Figure 4. This rearrangement is derived by the rotation of the H3R129’ side chain, that comes to interact with the modification site (OCS) and pushes the H3*α*2 helices away from each other. This also results in the disruption of the very stable H113-D123’ and H113’-D123 interactions that maintain the four-helix bundle in the canonical nucleosome. Long-range effects are observed, with a weakening of the H3-H4 communication pathways and hubs within a radius of *∼*15 Å from the OCS site, and the perturbation of the hydrogen bonds network especially near H3/H3’ C-termini through the disruption of interaction with H2A and H4 - see Figure S6. However, upon OCS the communication pathways are more intense and hubs get denser in both H3’ *α*2 and H2A *α*2, with larger communication blocks over the nucleosome structure. These observations suggest that the S-sulfonylation might destabilize the H3-H3 interface while stabilizing each separate dimer, which could be important for promoting the nucleosome disassembly process.

Besides the histone core architecture, H3C110 S-sulfonylation also destabilizes DNA near the dyad axis. The latter is normally the most strongly positioned part of the DNA in the nucleosome, and is the last region of DNA to be detached from the (H3-H4)_2_ tetramer during the nucleosome disassembly (45). We showed here that the DNA section between SHL0 and SHL-1 exhibits an increased flexibility upon histone H3 S-sulfonylation, not directly above the modification site but in symmetric with respect to the dyad axis - see Figure 6-a and b. This effect seems to results from very fine structural effects linked to the OCS-induced H3 *α*2 helices displacement. While the DNA-protein contacts at the histone core lateral surface exhibit the same patterns in the canonical and hyperoxidized nucleosome structures, the distribution of hydrogen bonds involving H3T118’ (SHL-1) and H3K115 (SHL0) are broader upon OCS, suggesting less stable interactions in these regions. Likewise, the base pairs structural parameters within the SHL+1/SHL-1 DNA section exhibit only weak deviations, yet they are mostly located in the SHL0/SHL-1 region which is also the one exhibiting an enhanced flexibility - see Figure 6-c and d. These observations suggest that S-sulfonylation destabilizes DNA at the dyad in an indirect way, through a subtle reorganization of the interaction network in the H3-H4 dimers. Of course, histone PTM rarely happen alone and oxidative stress conditions might result in the modification of other residues besides H3C110, which could have a synergistic effect on the nucleosome dynamics. Experimental investigations are needed to confirm the role of this modification and its interplay with other oxidative PTM in the nucleosome disassembly process upon oxidative stress.

Noteworthy, experimental data about histone H3 S-sulfonylation formation and function are scarce, and it is not known if this modification occurs before or after the assembly of the nucleosome. H3C110 is highly buried within the nucleosome structure, but it is known that S-glutathionylation (a much more bulky PTM) can be formed in the nucleosome context. Hence, it is reasonable to hypothesize that cysteine reaction with reactive oxygen species (ROS) could allow the formation of S-sulfonylated C110 in the nucleosome, which might play a different role than in free histones (e.g., nucleosome destabilization vs hindering of histone chaperone binding). Experimental investigations could shed light on this point.

## CONCLUSION

Among the variety of epigenetic marks regulating DNA compaction, oxidative PTM are very little studied. Resorting to extensive molecular dynamics simulations and in-depth structural analysis, we described the intrinsic communication networks within the canonical nucleosome and how they are re-shaped by S-sulfonylation of histone H3. This PTM results from the hyperoxidation of H3C110 upon oxidative stress conditions. We showed that this oxidative PTM induces subtle structural rearrangements that not only impact the histone core architecture, but also destabilize histone-DNA interactions near the dyad axis. These investigations constitute the first extensive insights into the alteration of nucleosome interaction pathways by a histone oxidative modification. Our results suggest that upon oxidative stress S-sulfonylation might contribute to the standard mechanisms promoting nucleosomal disassembly, and they provide an atomic-scale description of the mechanistic details underlying this process. This work provides an important computational framework for the study of other PTM, disease-related mutations, and histone variants effects onto the nucleosome architecture, that will shed light onto the finely-tuned mechanisms underlying DNA compaction regulation.

## Supporting information

Sup_Mat

## ACKNOWLEDGEMENTS

This work was granted access to the HPC resources IDRIS under the allocation 2022-A0120713412 made by GENCI. E.B thanks the Explor computing center for computational resources. E.B is grateful to Dr. Tao Jiang and Prof. Elise Dumont for providing flexibility analysis scripts.

## REFERENCES

1. Davey, C. A., Sargent, D. F., Luger, K., Maeder, A. W., and Richmond, T. J. (2002) Solvent mediated interactions in the structure of the nucleosome core particle at 1.9 Å resolution. Journal of molecular biology, 319(5), 1097–1113.

2. Luger, K., Maäder, A. W., Richmond, R. K., Sargent, D. F., and Richmond, T. J. (1997) Crystal structure of the nucleosome core particle at 2.8 resolution. Nature, 389(6648), 251–260.

3. Garcia-Saez, I., Menoni, H., Boopathi, R., Shukla, M. S., Soueidan, L., Noirclerc-Savoye, M., Le Roy, A., Skoufias, D. A., Bednar, J., Hamiche, A., et al. (2018) Structure of an H1-bound 6-nucleosome array reveals an untwisted two-start chromatin fiber conformation. Molecular cell, 72(5), 902–915.

4. Henikoff, S. (2008) Nucleosome destabilization in the epigenetic regulation of gene expression. Nature Reviews Genetics, 9(1), 15–26.

5. Armeev, G. A., Gribkova, A. K., Pospelova, I., Komarova, G. A., and Shaytan, A. K. (2019) Linking chromatin composition and structural dynamics at the nucleosome level. Current Opinion in Structural Biology, 56, 46–55.

6. Bignon, E., Allega, M. F., Lucchetta, M., Tiberti, M., and Papaleo, E. (2018) Computational structural biology of S-nitrosylation of cancer targets. Frontiers in oncology, 8, 272.

7. García-Giménez, J.-L., Garcés, C., Romá-Mateo, C., and Pallardó, F. V. (2021) Oxidative stress-mediated alterations in histone post-translational modifications. Free Radical Biology and Medicine, 170, 6–18.

8. García-Giménez, J. L., Romá-Mateo, C., and Pallardó, F. V. (2019) Oxidative post-translational modifications in histones. Biofactors, 45(5), 641–650.

9. Sha, Y. and Marshall, H. E. (2012) S-nitrosylation in the regulation of gene transcription. Biochimica et Biophysica Acta (BBA)-General Subjects, 1820(6), 701–711.

10. Marinho, H. S., Real, C., Cyrne, L., Soares, H., and Antunes, F. (2014) Hydrogen peroxide sensing, signaling and regulation of transcription factors. Redox biology, 2, 535–562.

11. Kim, Y.-J., Kim, D., Illuzzi, J. L., Delaplane, S., Su, D., Bernier, M., Gross, M. L., Georgiadis, M. M., and Wilson III, D. M. (2011) S-glutathionylation of cysteine 99 in the APE1 protein impairs abasic endonuclease activity. Journal of molecular biology, 414(3), 313–326.

12. Tang, C.-H., Wei, W., and Liu, L. (2012) Regulation of DNA repair by S-nitrosylation. Biochimica et biophysica acta (BBA)-general subjects, 1820(6), 730–735.

13. García-Giménez, J. L., Olaso, G., Hake, S. B., Boänisch, C., Wiedemann, S. M., Markovic, J., Dasi, F., Gimeno, A., Pérez-Quilis, C., Palacios, O., et al. (2013) Histone h3 glutathionylation in proliferating mammalian cells destabilizes nucleosomal structure. Antioxidants & redox signaling, 19(12), 1305–1320.

14. Simon, M., North, J. A., Shimko, J. C., Forties, R. A., Ferdinand, M. B., Manohar, M., Zhang, M., Fishel, R., Ottesen, J. J., and Poirier, M. G. (2011) Histone fold modifications control nucleosome unwrapping and disassembly. Proceedings of the National Academy of Sciences, 108(31), 12711–12716.

15. Bowman, G. D. and Poirier, M. G. (2014) Post-translational modifications of histones that influence nucleosome dynamics. Chemical reviews, 115(6), 2274–2295.

16. Chatterjee, N., North, J. A., Dechassa, M. L., Manohar, M., Prasad, R., Luger, K., Ottesen, J. J., Poirier, M. G., and Bartholomew, B. (2015) Histone acetylation near the nucleosome dyad axis enhances nucleosome disassembly by RSC and SWI/SNF. Molecular and cellular biology, 35(23), 4083–4092.

17. North, J. A., Šimon, M., Ferdinand, M. B., Shoffner, M. A., Picking, J. W., Howard, C. J., Mooney, A. M., van Noort, J., Poirier, M. G., and Ottesen, J. J. (2014) Histone H3 phosphorylation near the nucleosome dyad alters chromatin structure. Nucleic acids research, 42(8), 4922–4933.

18. Tropberger, P., Pott, S., Keller, C., Kamieniarz-Gdula, K., Caron, M., Richter, F., Li, G., Mittler, G., Liu, E. T., Buähler, M., et al. (2013) Regulation of transcription through acetylation of H3K122 on the lateral surface of the histone octamer. Cell, 152(4), 859–872.

19. Bowerman, S. and Wereszczynski, J. (2016) Effects of MacroH2A and H2A. Z on nucleosome dynamics as elucidated by molecular dynamics simulations. Biophysical journal, 110(2), 327–337.

20. Phillips, J. C., Hardy, D. J., Maia, J. D., Stone, J. E., Ribeiro, J. V., Bernardi, R. C., Buch, R., Fiorin, G., Hénin, J., Jiang, W., et al. (2020) Scalable molecular dynamics on CPU and GPU architectures with NAMD. J. Chem. Phys., 153(4), 044130.

21. Case, D., Belfon, K., Ben-Shalom, I., Brozell, S., Cerutti, D., Cheatham III, T., Cruzeiro, V., Darden, T., Duke, R., Giambasu, G., Gilson, M., Gohlke, H., Goetz, A., Harris, R., Izadi, S., Izmailov, S., Kasavajhala, K., Kovalenko, A., Krasny, R., Kurtzman, T., Lee, T., LeGrand, S., Li, P., Lin, C., Liu, J., Luchko, T., Luo, R., Man, V., Merz, K., Miao, Y., Mikhailovskii, O., Monard, G., Nguyen, H., Onufriev, A., Pan, ÅF., Pantano, S., Qi, R., Roe, D., Roitberg, A., Sagui, C., Schott-Verdugo, S., Shen, J., Simmerling, C., Skrynnikov, N., Smith, J., Swails, J., Walker, R., Wang, J., Wilson, L., Wolf, R., Wu, X., Xiong, Y., Xue, Y., York, D., and Kollman, P. (2020) AMBER 2020. University of California, San Francisco.

22. Karami, Y., Bitard-Feildel, T., Laine, E., and Carbone, A. (2018) “Infostery” analysis of short molecular dynamics simulations identifies highly sensitive residues and predicts deleterious mutations. Scientific reports, 8(1), 16126.

23. Lavery, R., Moakher, M., Maddocks, J. H., Petkeviciute, D., and Zakrzewska, K. (2009) Conformational analysis of nucleic acids revisited: Curves+. Nucleic Acids Research, 37(17), 5917–5929.

24. Humphrey, W., Dalke, A., and Schulten, K. (1996) VMD: visual molecular dynamics. J. Mol. Graph., 14(1), 33–38.

25. Schroädinger, LLC (November, 2015) The PyMOL Molecular Graphics System, Version 1.8.

26. Maier, J. A., Martinez, C., Kasavajhala, K., Wickstrom, L., Hauser, K. E., and Simmerling, C. (2015) ff14SB: improving the accuracy of protein side chain and backbone parameters from ff99SB. Journal of chemical theory and computation, 11(8), 3696–3713.

27. Ivani, I., Dans, P. D., Noy, A., Perez, A., Faustino, I., Hospital, A., Walther, J., Andrio, P., Goni, R., Balaceanu, A., Portella, G., Battistini, F., Gelpí, J. L., González, C., Vendruscolo, M., Laughton, C. A., Harris, S. A., Case, D. A.,, and Orozco, M. (2016) Parmbsc1: a refined force field for DNA simulations. Nature Methods, 38(13), 55–58.

28. Yoo, J. and Aksimentiev, A. (2018) New tricks for old dogs: improving the accuracy of biomolecular force fields by pair-specific corrections to non-bonded interactions. Physical Chemistry Chemical Physics, 20(13), 8432–8449.

29. Bhatt, M. R. and Zondlo, N. J. (2023) Synthesis and conformational preferences of peptides and proteins with cysteine sulfonic acid. Organic & Biomolecular Chemistry, 21(13), 2779–2800.

30. Hopkins, C. W., Le Grand, S., Walker, R. C., and Roitberg, A. E. (apr, 2015) Long-time-step molecular dynamics through hydrogen mass repartitioning. Journal of Chemical Theory and Computation, 11(4), 1864–1874.

31. Miyamoto, S. and Kollman, P. A. (oct, 1992) Settle: An analytical version of the SHAKE and RATTLE algorithm for rigid water models. Journal of Computational Chemistry, 13(8), 952–962.

32. Darden, T., York, D., and Pedersen, L. (1993) Particle mesh Ewald: An N log (N) method for Ewald sums in large systems. The Journal of chemical physics, 98(12), 10089–10092.

33. Smirnova, E., Bignon, E., Schultz, P., Papai, G., and Ben-Shem, A. (2023) Binding to nucleosome poises SIRT6 for histone H3 de-acetylation. eLife, 12, RP87989.

34. Karami, Y., Laine, E., and Carbone, A. (2016) Dissecting protein architecture with communication blocks and communicating segment pairs. BMC Bioinformatics, 17 Suppl 2(Suppl 2), 13.

35. McDonald, I. K. and Thornton, J. M. (1994) Satisfying hydrogen bonding potential in proteins. Journal of molecular biology, 238(5), 777–793.

36. Bignon, E., Gillet, N., Chan, C.-H., Jiang, T., Monari, A., and Dumont, E. (2021) Recognition of a tandem lesion by DNA bacterial formamidopyrimidine glycosylases explored combining molecular dynamics and machine learning. Computational and structural biotechnology journal, 19, 2861–2869.

37. Bignon, E., Claerbout, V. E., Jiang, T., Morell, C., Gillet, N., and Dumont, E. (2020) Nucleosomal embedding reshapes the dynamics of abasic sites. Scientific reports, 10(1), 17314.

38. Bignon, E., Gillet, N., Jiang, T., Morell, C., and Dumont, E. (2021) A dynamic view of the interaction of histone tails with clustered abasic sites in a nucleosome core particle. The Journal of Physical Chemistry Letters, 12(25), 6014–6019.

39. Cutter, A. R. and Hayes, J. J. (2015) A brief review of nucleosome structure. FEBS letters, 589(20), 2914–2922.

40. Souza, V. P., Ikegami, C. M., Arantes, G. M., and Marana, S. R. (2018) Mutations close to a hub residue affect the distant active site of a GH1 β-glucosidase. Plos one, 13(6), e0198696.

41. Banks, D. D. and Gloss, L. M. (2003) Equilibrium Folding of the Core Histones: the H3-H4 Tetramer Is Less Stable than the H2A-H2B Dimer. Biochemistry, 42(22), 6827–6839.

42. Xu, Y.-M., Du, J.-Y., and Lau, A. T. (2014) Posttranslational modifications of human histone H3: an update. Proteomics, 14(17-18), 2047–2060.

43. Nacev, B. A., Feng, L., Bagert, J. D., Lemiesz, A. E., Gao, J., Soshnev, A. A., Kundra, R., Schultz, N., Muir, T. W., and Allis, C. D. (2019) The expanding landscape of ‘oncohistone’mutations in human cancers. Nature, 567(7749), 473–478.

44. Tessarz, P. and Kouzarides, T. (2014) Histone core modifications regulating nucleosome structure and dynamics. Nature reviews Molecular cell biology, 15(11), 703–708.

45. Gansen, A., Valeri, A., Hauger, F., Felekyan, S., Kalinin, S., Tóth, K., Langowski, J., and Seidel, C. A. (2009) Nucleosome disassembly intermediates characterized by single-molecule FRET. Proceedings of the National Academy of Sciences, 106(36), 15308–15313.

