## Supplementary material for "Cysteine hyperoxidation rewires communication pathways in the nucleosome and destabilizes the dyad": Sup_Mat

### Supporting Information for: Cysteine hyperoxidation rewires communication pathways in the nucleosome and destabilizes the dyad

Yasaman Karami† and Emmanuelle Bignon*,‡

†*Université de Lorraine, CNRS, Inria, LORIA, F-54000 Nancy, France*

‡*Université de Lorraine and CNRS, LPCT UMR 7019, F-54000 Nancy, France*

Table S1 – Block size for each system. The number residues within each communication block are reported for the all the identified blocks.

| Block | 601 Widom | 1KX5 | 1KX5 + OCS |
|---|---|---|---|
| 1 | 12 | 13 | 14 |
| 2 | 57 | 7 | 57 |
| 3 | 58 | 37 | 62 |
| 4 | 136 | 10 | 139 |
| 5 | 13 | 44 | 12 |
| 6 | 11 | 4 | 123 |
| 7 | 36 | 15 | 4 |
| 8 | 64 | 112 | 14 |
| 9 | 131 | 13 | 109 |
| 10 | - | 12 | - |
| 11 | - | 4 | - |
| 12 | - | 32 | - |
| 13 | - | 56 | - |
| 14 | - | 13 | - |
| 15 | - | 115 | - |

Table S2 – List of the monitored interactions between the lateral surface of the nucleosome and DNA. Amino acids from the second copy of each histone is marked by an apostrophe. Interactions found near the dyad (i.e., between SHL-1 and SHL+1) are marked by a star.

| | | | |
|---|---|---|---|
| **H3** | | | |
| H3Y41-dA231* | H3Y41'-dA84* | H3R42-dG145 | H3R42'-dG292 |
| H4K44'-dT217* | H3T45-dG145 | H3T45'-dG292 | H3R49'-dT8 |
| H3K56-dC156 | H3K56'-dC9 | H3K64-dT239 | H3K64'-dT92 |
| H3R83-dC50 | H3R83'-dG101 | H3S86-dC50 | H3K115-dA220* |
| H3K115'-dA73* | H3T118-dG218* | H3T118'-dG71* | H3K122-dG72* |
| H3K122'-dG219* | | | |
| **H4** | | | |
| H4T30'-dA208 | H4R35-dG229* | H4R39'-dG82* | H4R45-dT228* |
| H4R45'-dT81* | H4S47-dT228* | H4S47'-dT81* | H4K77-dA41 |
| H4K77'-dA188 | H4K79-dC49 | H4K79'-dC102 | H4T80-dC249 |
| **H2A** | | | |
| H2AR32'-dA177 | H2A42-dT38 | H2A42'-dG186 | H2AT76-dC280 |
| H2AT76'-dC133 | H2AR77-dA19 | H2AR77'-dC133 | |
| **H2B** | | | |
| H2BY37-dG269 | H2BS53-dG40 | H2BS53'-dA166 | H2BR83-dG40 |
| H2BR83'-dA188 | H2BS84-dG40 | H2BT85-dG40 | H2BT85'-dG187 |

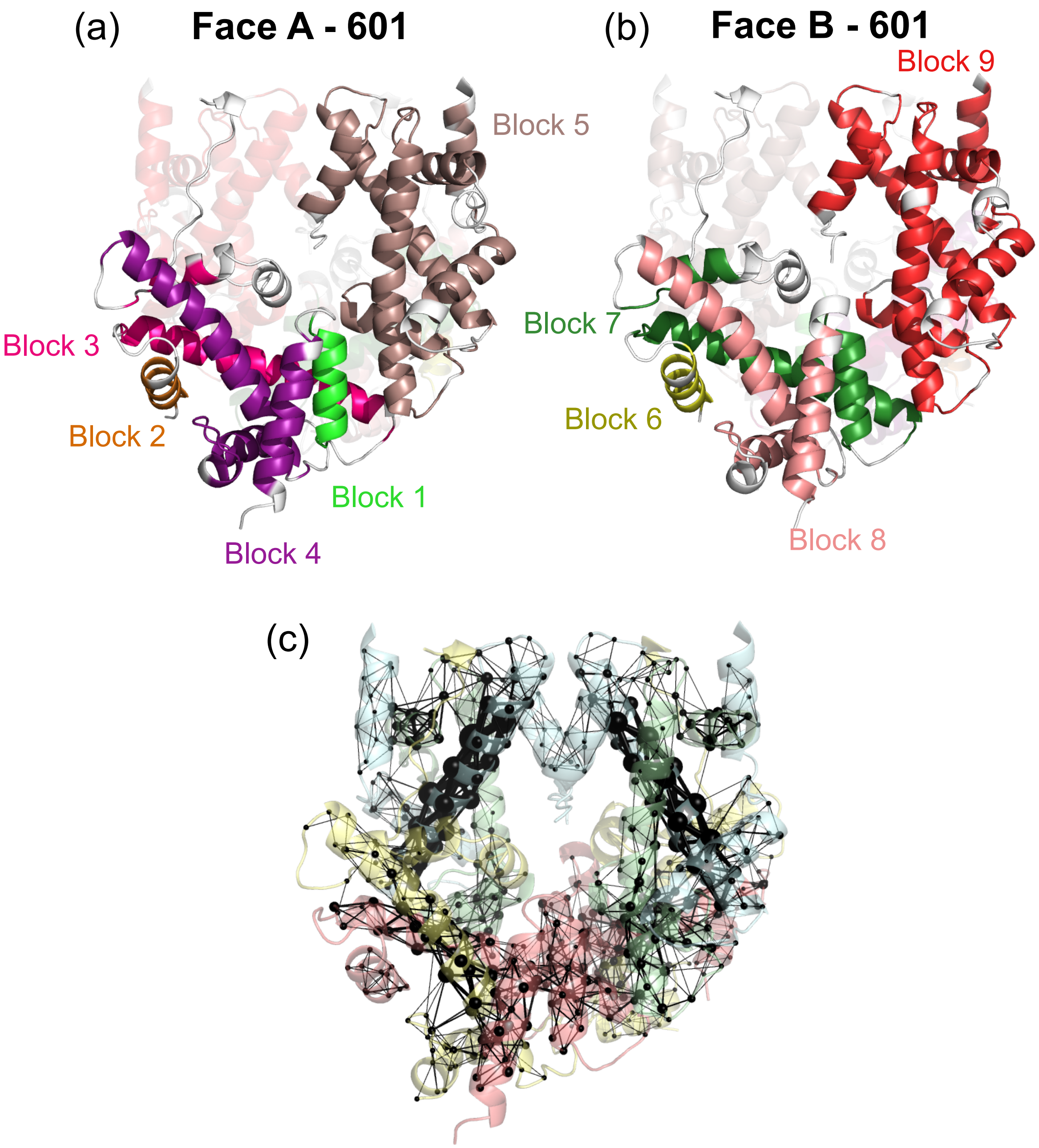


Figure S1 – Communication blocks in the 601 nucleosome, viewed from **(a)** face A and **(b)** face B, and **(c)** corresponding communication pathways.

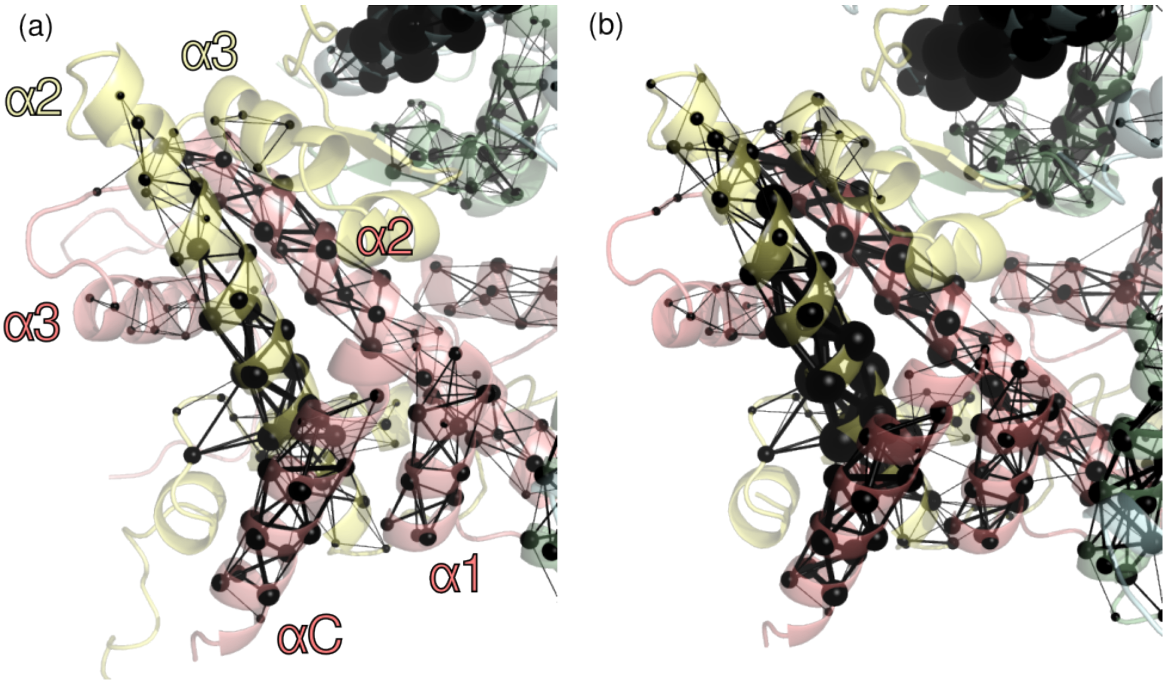


Figure S2 – Communication pathways and hubs within the H2A-H2B dimer, without S-sulfonylation **(a)** or with it **(b)**.

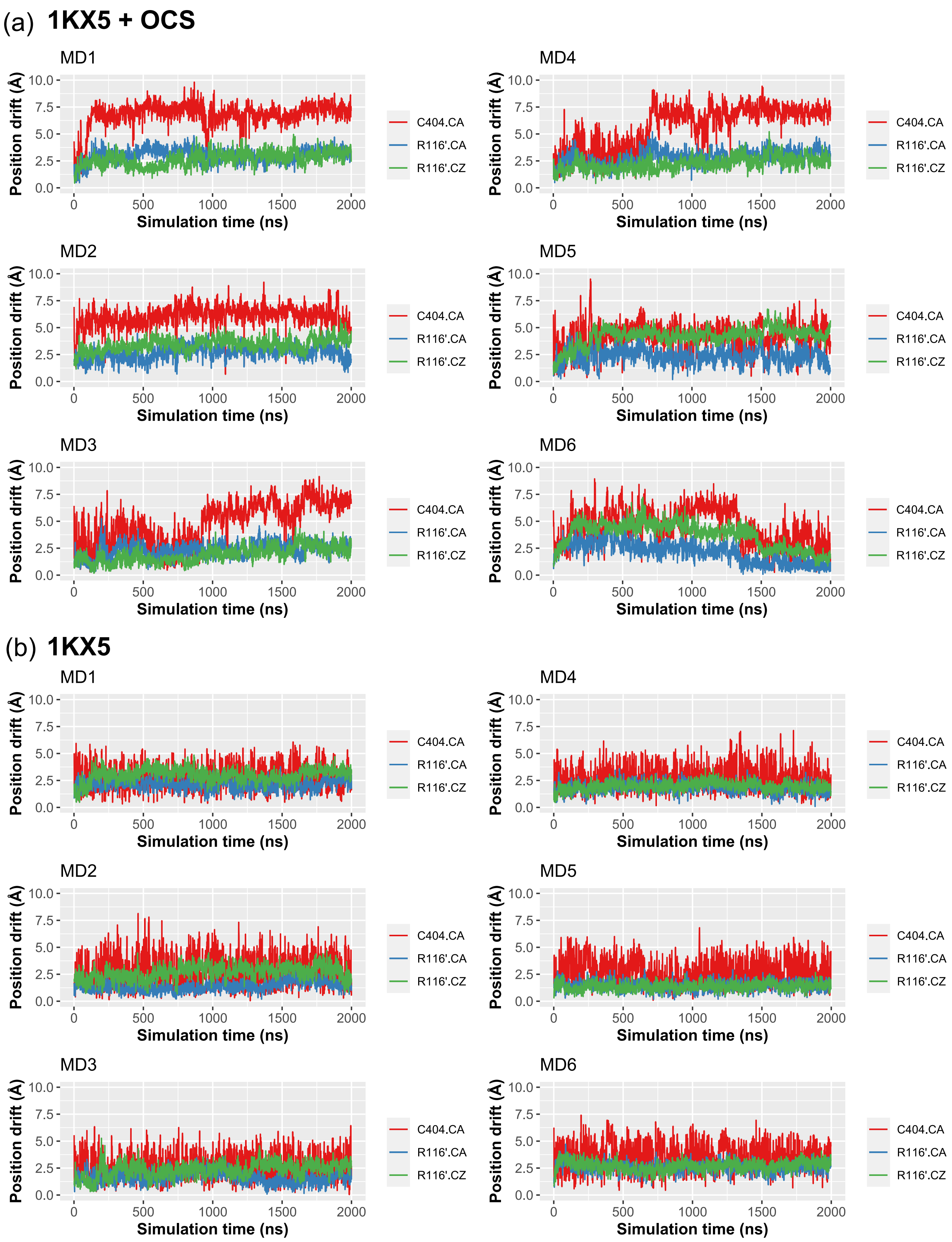


Figure S3 – Evolution of the positional drift of C110 CA atom and R116’ CA and CZ atoms in the 6 replicates of simulations **(a)** with OCS and **(b)** without it.

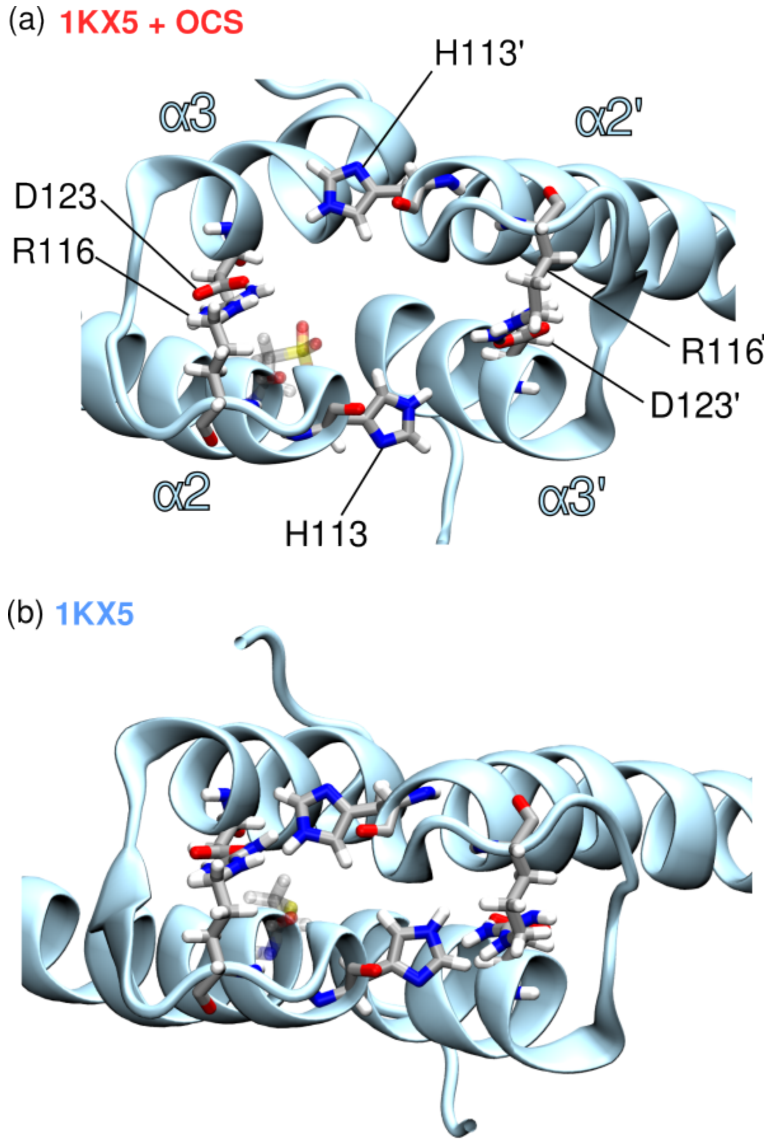


Figure S4 – Top view of the H3-H3' four-helix bundle, highlighting the interaction network involving H113('), D123(') and R116(') residues, in simulations **(a)** with OCS and **(b)** without it.

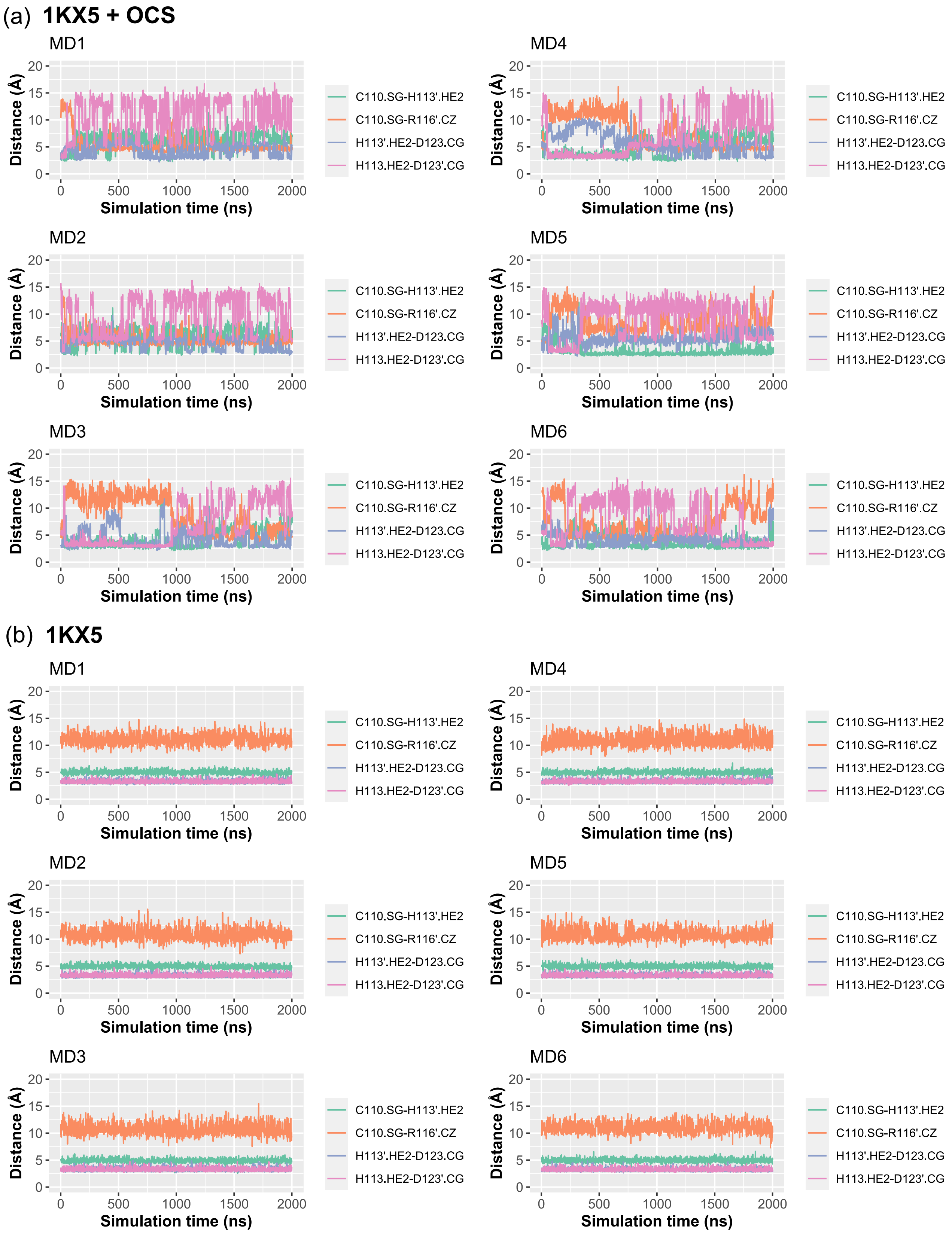


Figure S5 – Evolution of the key-interactions within the H3-H3 four-helix bundle in the 6 replicates of simulations **(a)** with OCS and **(b)** without it.

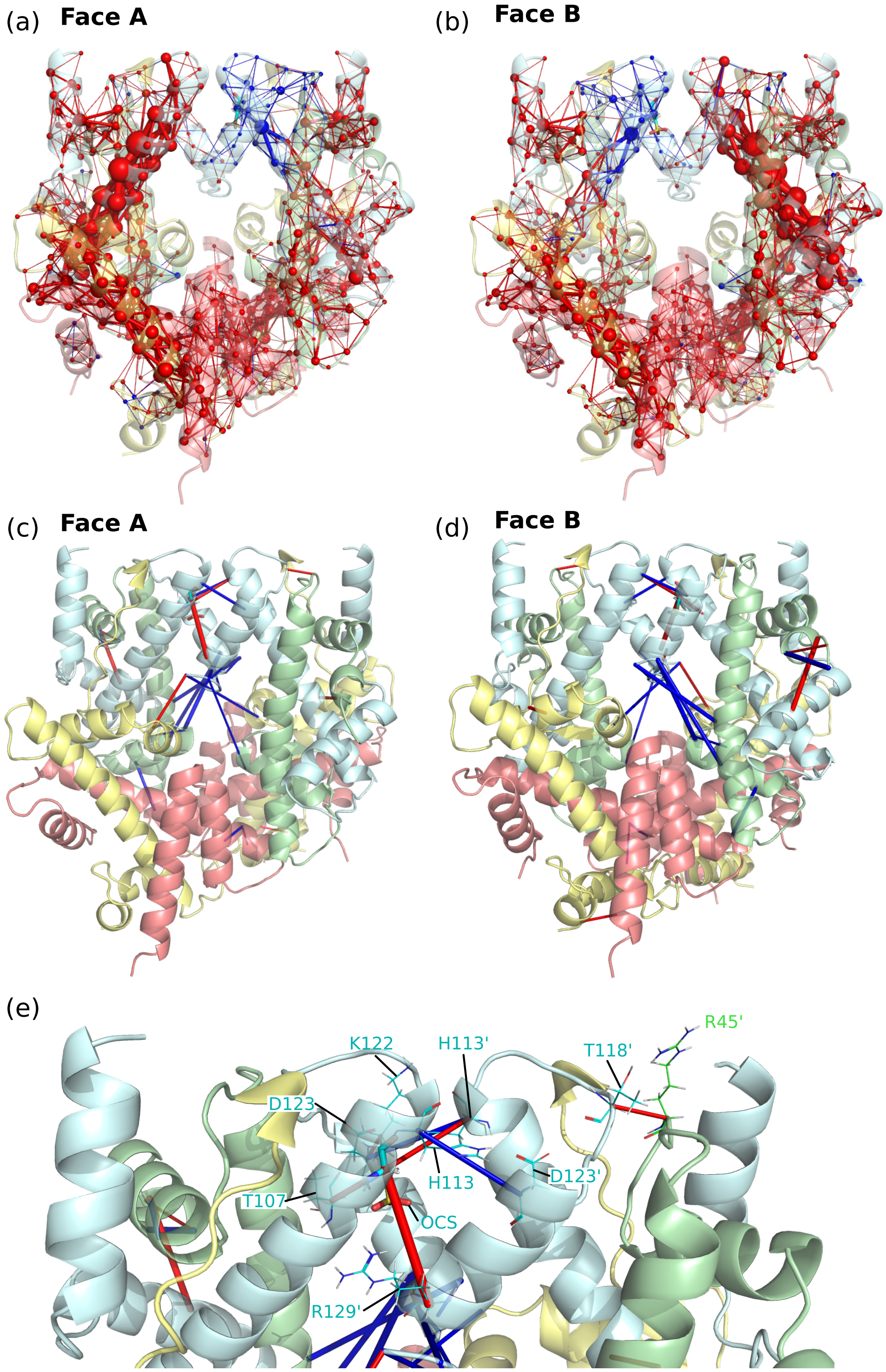


Figure S6 – **(a)** Perturbation of the communication pathways upon S-sulfonylation. Blue spheres/sticks represent lost hubs/pathways, and red ones the gained hubs/pathways, showed for face A and **(b)** face B. **(c)** Hydrogen bonds network changes upon S-sulfonylation (blue=loss, red=gain), viewed from face A and **(d)** face B. **(e)** Zoom on the hydrogen bond loss and gain in the four-helix bundle corresponding to the H3-H3 contact region.

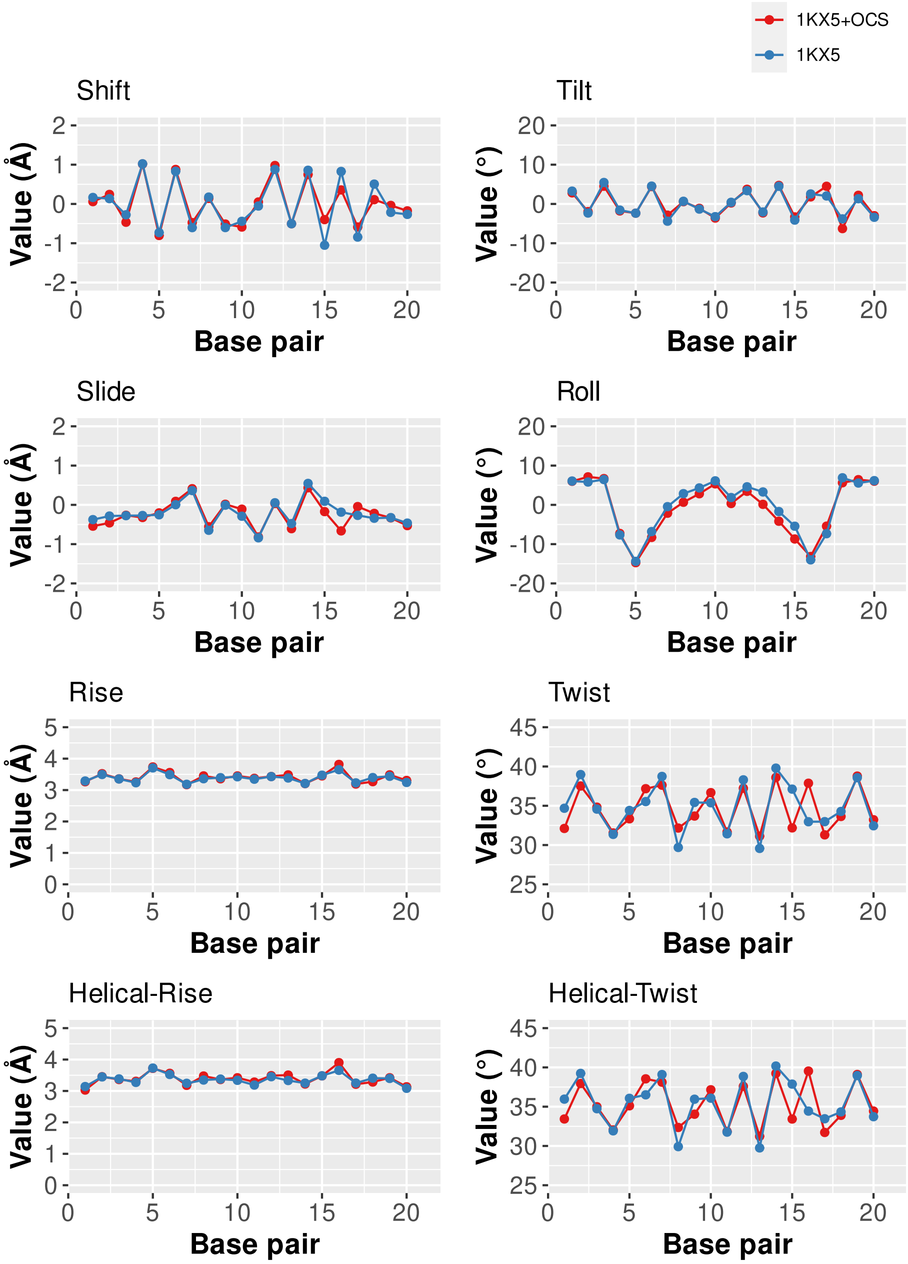


Figure S7 – Inter-base pair parameters in the SHL-1/SHL+1 section, for the canonical system (1KX5, blue) and the S-sulfonylated system (1KX5+OCS, red).

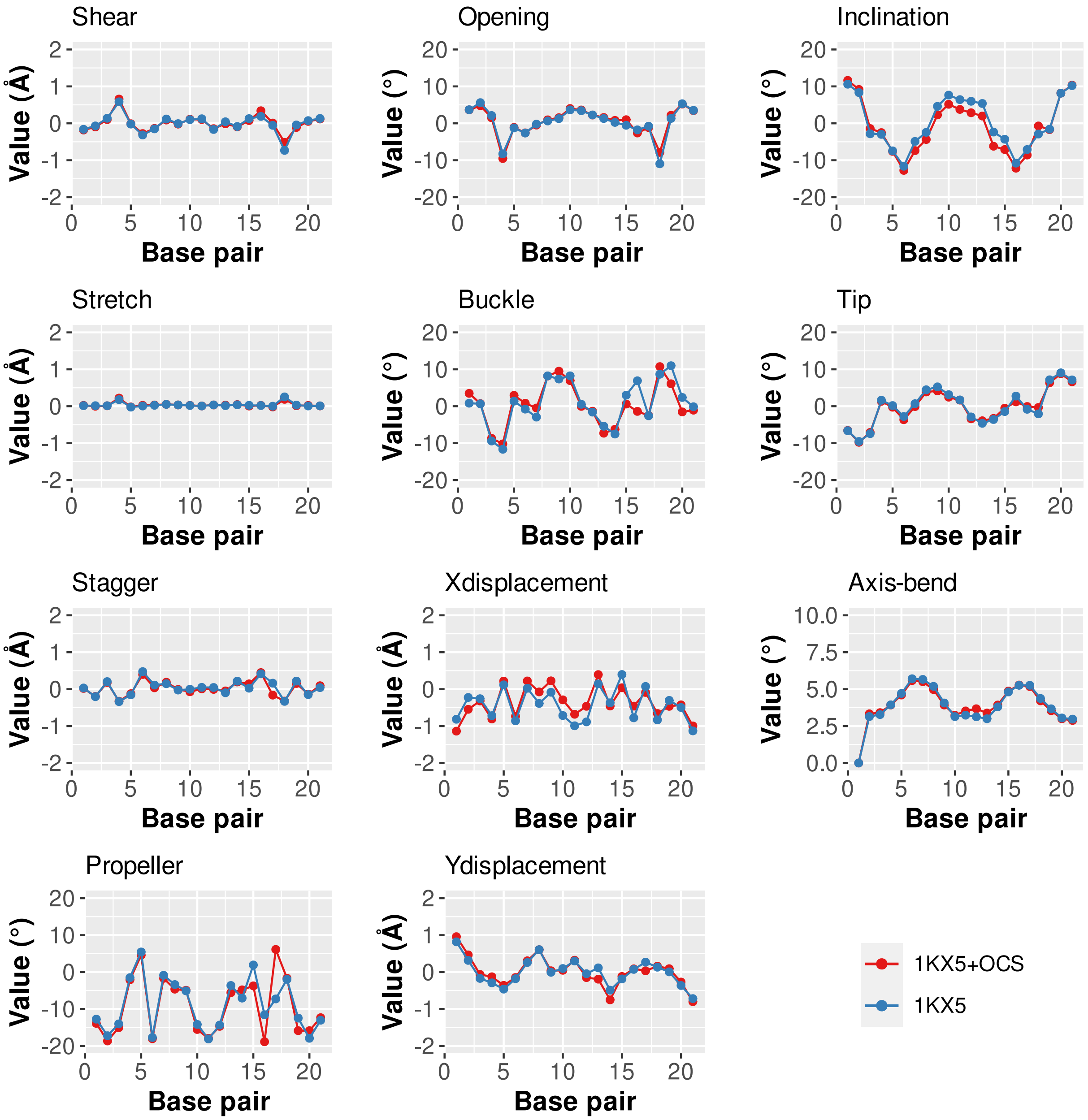


Figure S8 – Intra-base pair and base pair axis parameters in the SHL-1/SHL+1 section, for the canonical system (1KX5, blue) and the S-sulfonylated system (1KX5+OCS, red).

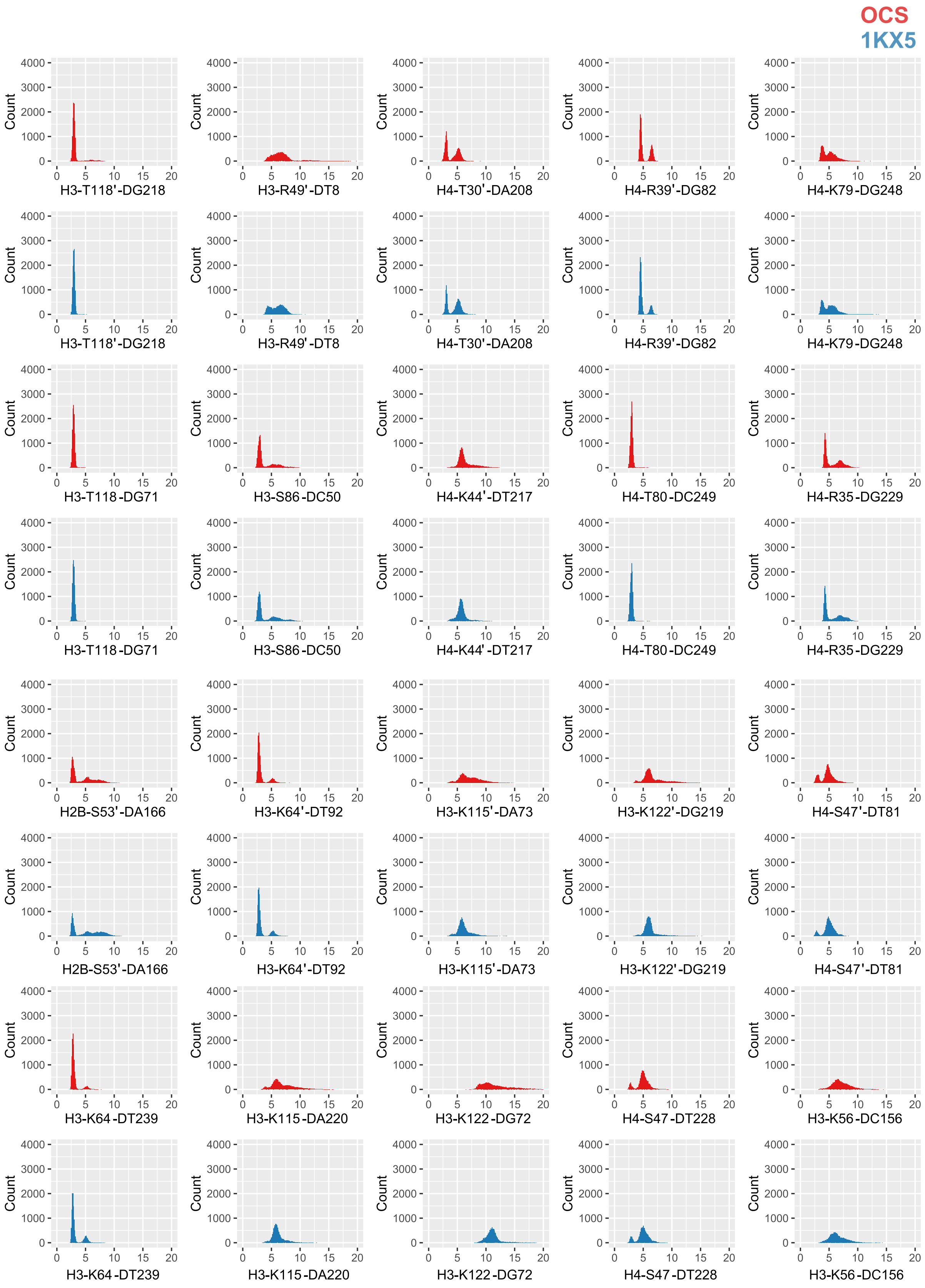


Figure S9 – Distribution of the distances corresponding to the histone core lateral surface interactions with DNA (part 1), as listed in Table S2. Values are given for simulations with (red) or without (blue) OCS.

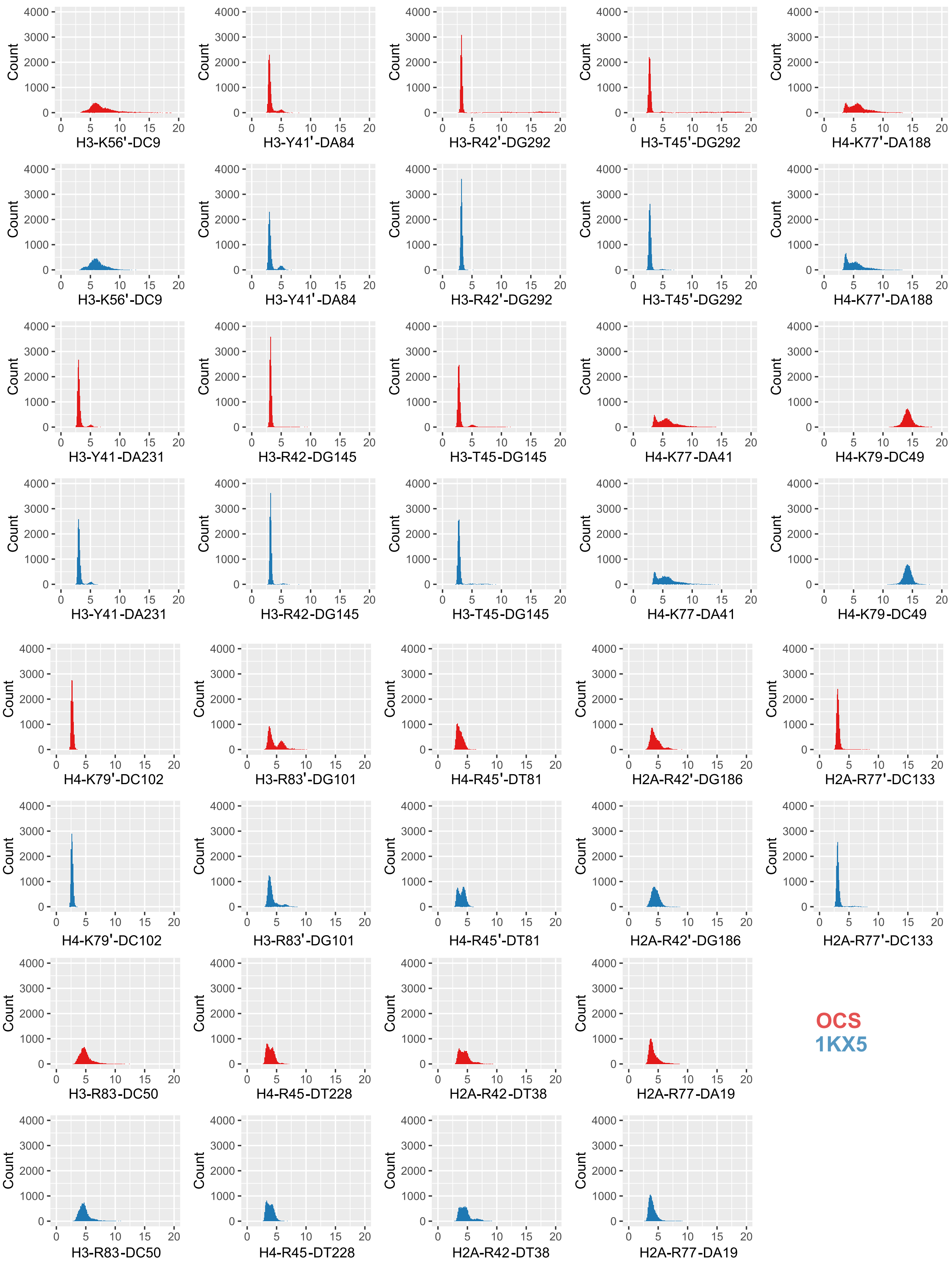


Figure S10 – Distribution of the distances corresponding to the histone core lateral surface interactions with DNA (part 2), as listed in Table S2. Values are given for simulations with (red) or without (blue) OCS.

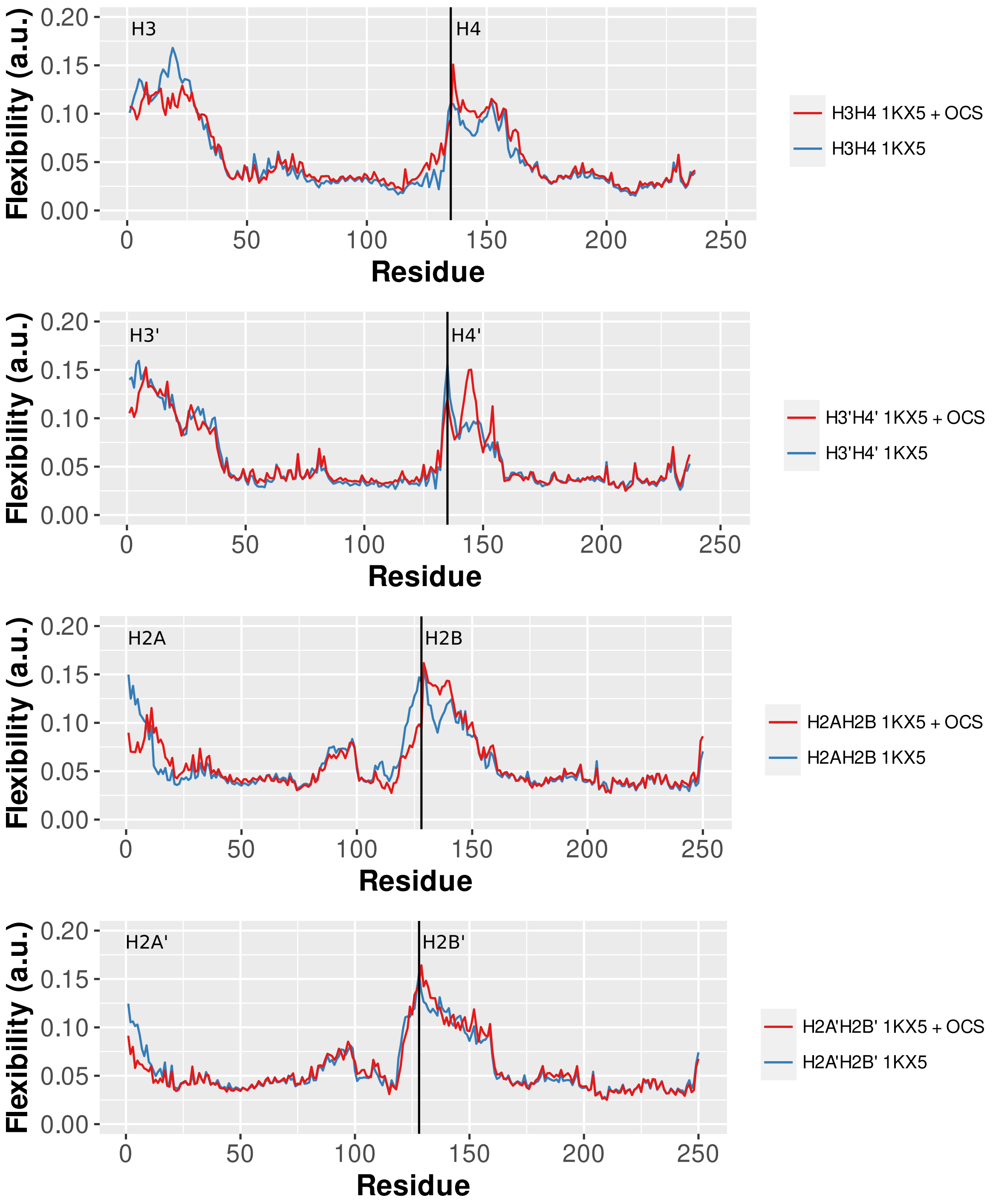


Figure S11 – Per residue flexibility in each dimer copy, for simulation of the canonical nucleosome with (1KX5 + OCS, red) and without (1KX5,blue) S-sulfonylation. Strong deviations are only found in the disordered N- and C-termini of the histones, suggesting that the modification do not impact the core flexbility. Of note, OCS is located in H3 at position 110.
